# Non-genetic adaptation by collective migration

**DOI:** 10.1101/2024.01.02.573956

**Authors:** Lam Vo, Fotios Avgidis, Henry H. Mattingly, Karah Edmonds, Isabel Burger, Ravi Balasubramanian, Thomas S. Shimizu, Barbara I. Kazmierczak, Thierry Emonet

## Abstract

Cell populations must adjust their phenotypic composition to adapt to changing environments. One adaptation strategy is to maintain distinct phenotypic subsets within the population and to modulate their relative abundances via gene regulation. Another strategy involves genetic mutations, which can be augmented by stress-response pathways. Here, we studied how a migrating bacterial population regulates its phenotypic distribution to traverse diverse environments. We generated isogenic *Escherichia coli* populations with varying distributions of swimming behaviors and observed their phenotype distributions during migration in liquid and porous environments. We found that the migrating populations became enriched with high-performing swimming phenotypes in each environment, allowing the populations to adapt without requiring mutations or gene regulation. This adaptation is dynamic and rapid, reversing in a few doubling times when migration ceases. By measuring the chemoreceptor abundance distributions during migration towards different attractants, we demonstrated that adaptation acts on multiple chemotaxis-related traits simultaneously. These measurements are consistent with a general mechanism in which adaptation results from a balance between cell growth generating diversity and collective migration eliminating under-performing phenotypes. Thus, collective migration enables cell populations with continuous, multi-dimensional phenotypes to flexibly and rapidly adapt their phenotypic composition to diverse environmental conditions.

**Significance statement:** Conventional cell adaptation mechanisms, like gene regulation and stochastic phenotypic switching, act swiftly but are limited to a few traits, while mutation-driven adaptations unfold slowly. By quantifying phenotypic diversity during bacterial collective migration, we discovered an adaptation mechanism that rapidly and reversibly adjusts multiple traits simultaneously. By balancing the generation of diversity through growth with the loss of phenotypes unable to keep up, this process tunes the phenotypic composition of migrating populations to the environments they traverse, without gene regulation or mutations. Given the prevalence of collective migration in microbes, cancers, and embryonic development, non-genetic adaptation through collective migration may be a universal mechanism for populations to navigate diverse environments, offering insights into broader applications across various fields.

## Introduction

Cell populations use various mechanisms to adapt their phenotypic distribution to new environments. One approach is to acquire mutations, such as in bacterial efflux pumps^1,2^ or the binding sites of EGFR^3^ in cancer cells, to evade stressors. These processes typically occur over tens of generations^4–6^. Another common approach is to regulate gene expression or switching between discrete phenotypic states^7,8^. A classic example of phenotypic switching is persister formation, where bacteria^9–11^ or cancer cells^12^ transition between antibiotic-resistant and chemotherapy-tolerant states^13,14^.

Here, we examined whether and how a migrating bacterial population regulates its phenotypic composition according to the environments it encounters. Groups of *Escherichia coli* cells collectively migrate by consuming attractant cues in their environment and chasing the traveling gradient that they create^15,16^. Individual cells with the same genotype exhibit varying tumble biases (TB) (**Fig. 1a**), chemoreceptor abundances (**Fig. 1b**), and other chemotaxis-relevant phenotypes, leading to differences in the speeds at which they climb the attractant gradients^17^. These cells can travel together by spatially organizing within the migrating group such that their gradient-climbing abilities match the local steepness of the attractant gradient^15^ (**Fig. 1c**). However, this spatial sorting mechanism is imperfect: cells at the back, or the low performers, slowly fall behind the migrating population because the attractant concentration falls below the detection limit of their receptors^15^. To maintain migration, the lost cells must be replaced through cell divisions^18,19^.

**Figure 1:**
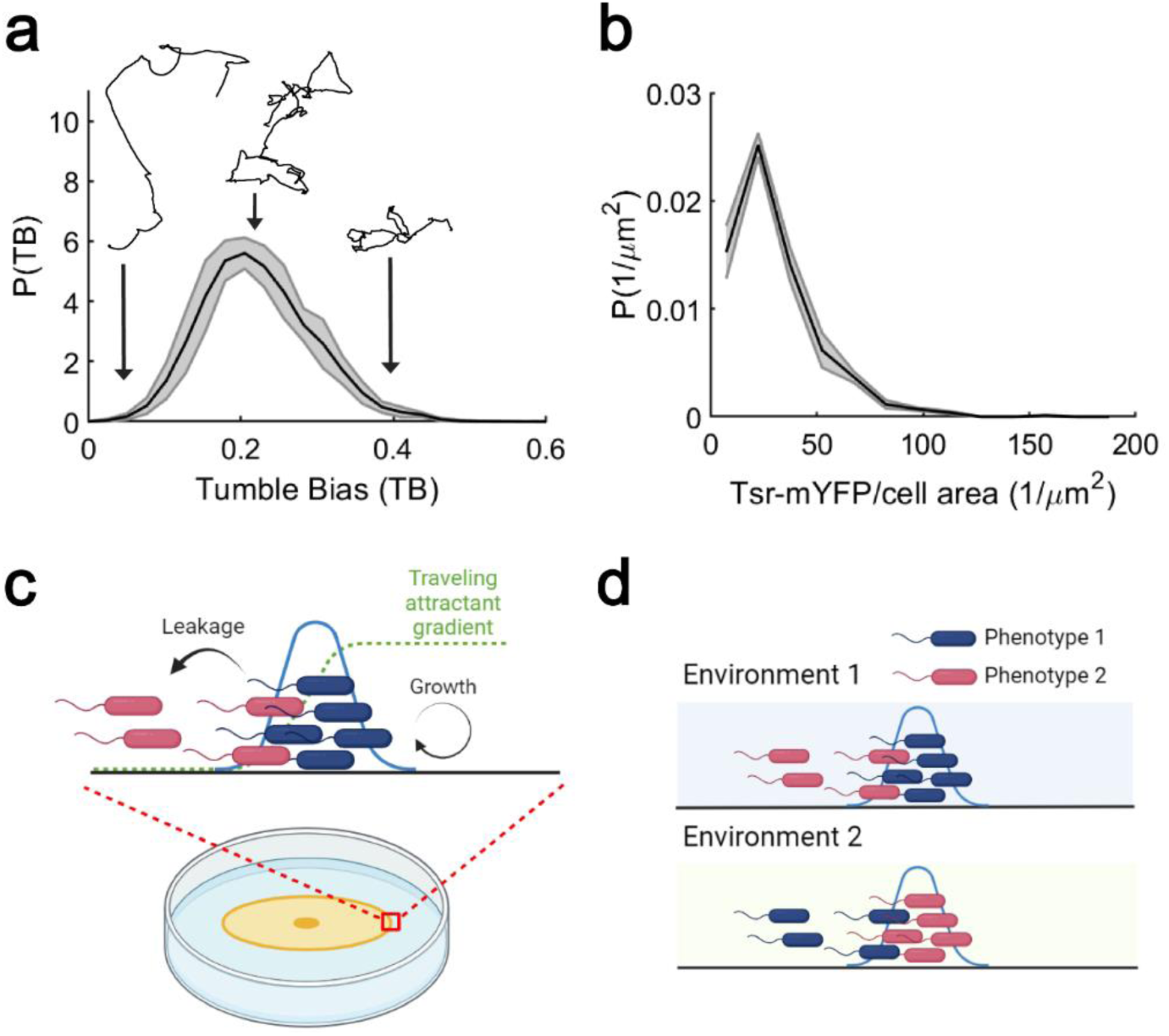
Isogenic populations of bacteria exhibit diversity in behavioral and sensory phenotypes while performing collective migration. **(a)** In an isogenic population of E. coli, there is diversity in swimming behaviors, which can be quantified, for example, by measuring tumble bias in E. coli RP437 (*N*_*rep*_ = 3, *N*_*cells*_ = 9,417). **(b)** There is also diversity in sensory ability, which correlates with variations in Tsr levels (as measured in E. coli MG1655, *N*_*rep*_ = 3, *N*_*cells*_ = 1,695). **(c)** Under nutrient-replete conditions, cells consume attractants and collectively migrate using the resulting traveling attractant gradients^21^. During this process, cells spatially organize based on their ability to climb the traveling gradients^15^. Cells with higher chemotactic performances lead at the front, while cells with lower chemotactic performances travel at the back and eventually fall behind the migrating group. In such nutrient-replete conditions, cells continue to divide and generate diversity by replenishing phenotypes. **(d)** Theory predicts^18^ that the balance between environment-dependent cell loss and growth enables the migrating isogenic population to dynamically tune its phenotypic composition to the environments it traverses.

A recent theoretical model proposed that a dynamic balance between loss of low-performing phenotypes due to migration and production of new phenotypes by growth might be sufficient to adapt the population’s composition to new environments^18^. These two processes would balance out such that the traveling population becomes enriched with “fitter” phenotypes—those with higher chemotactic performance—increasing its expansion speed^15,18^. Since the demands of migration are different in different environments, which phenotypes are enriched should depend on the environment the population traverses^18^ (**Fig. 1d**). This proposed mechanism does not require environment-dependent mutations or gene regulation. However, this model has never been experimentally tested.

Here, we experimentally demonstrate this novel adaptation mechanism. By measuring the distribution of swimming behaviors and chemoreceptor abundances during migration in distinct environments, we observed that migrating *E. coli* populations tune their own distributions of chemotactic behaviors, without relying on gene regulation or mutations. Adaptation relies on the balance between the differential loss of low-performing phenotypes and the generation of diversity by growth during migration in each environment. The relaxation from the adapted distribution to the standing batch distribution occurs in approximately two generations. More broadly, migration and other collective behaviors when combined with growth may generally provide rapid and flexible ways for diverse populations to adapt to changing conditions.

## Results

### Collective migration tunes the population’s distribution of swimming phenotypes

First, we investigated whether the TB distribution of an *E. coli* population can be adapted during collective migration with growth. Given that some swimming phenotypes are better at navigating than others^15,17^, we expected the TB distribution of a population, regardless of its starting point, to enrich for higher performing TBs during collective migration. To test this hypothesis, we compared the TB distributions of populations grown to exponential phase in batch cultures to those of populations migrating at constant speed on agarose swim plates containing excess nutrients. Excess nutrients ensure that cells in the migrating front are experiencing exponential growth ^20,21^.

Populations with different batch TB distributions were generated by controlling the expression levels of the phosphatase CheZ using an inducible promoter in cells lacking the endogenous *cheZ* gene, as previously described^15,22^ (**Methods**). Growing this strain with a high concentration (16 ng/mL) or low concentration (2 ng/mL) of the inducer anhydrotetracycline (aTc) resulted in a population with low average TB or high average TB, respectively (**Fig. S1**). The TB distribution of wild-type (WT) *E. coli* falls between these two extremes (**Fig. 1a**).

To propagate migrating waves of growing bacteria, we utilized the classic swim plate assay, in which cells migrate through a fibrous mesh of 0.14% w/v semisolid agarose supplemented with excess nutrients^16,23^. We inoculated *E. coli* populations with different batch TB distributions at the center of agarose swim plates, let them migrate for 15 hours, collected cells at the edge of the expanding colonies (“wave”), and measured their TB distributions by transferring the cells to a liquid buffer and tracking single cells under a microscope. In parallel, we grew these same populations with identical inducer concentrations in shaking flasks (“batch”) until exponential growth phase and measured their TB distributions (**Fig. 2a**).

**Figure 2.**
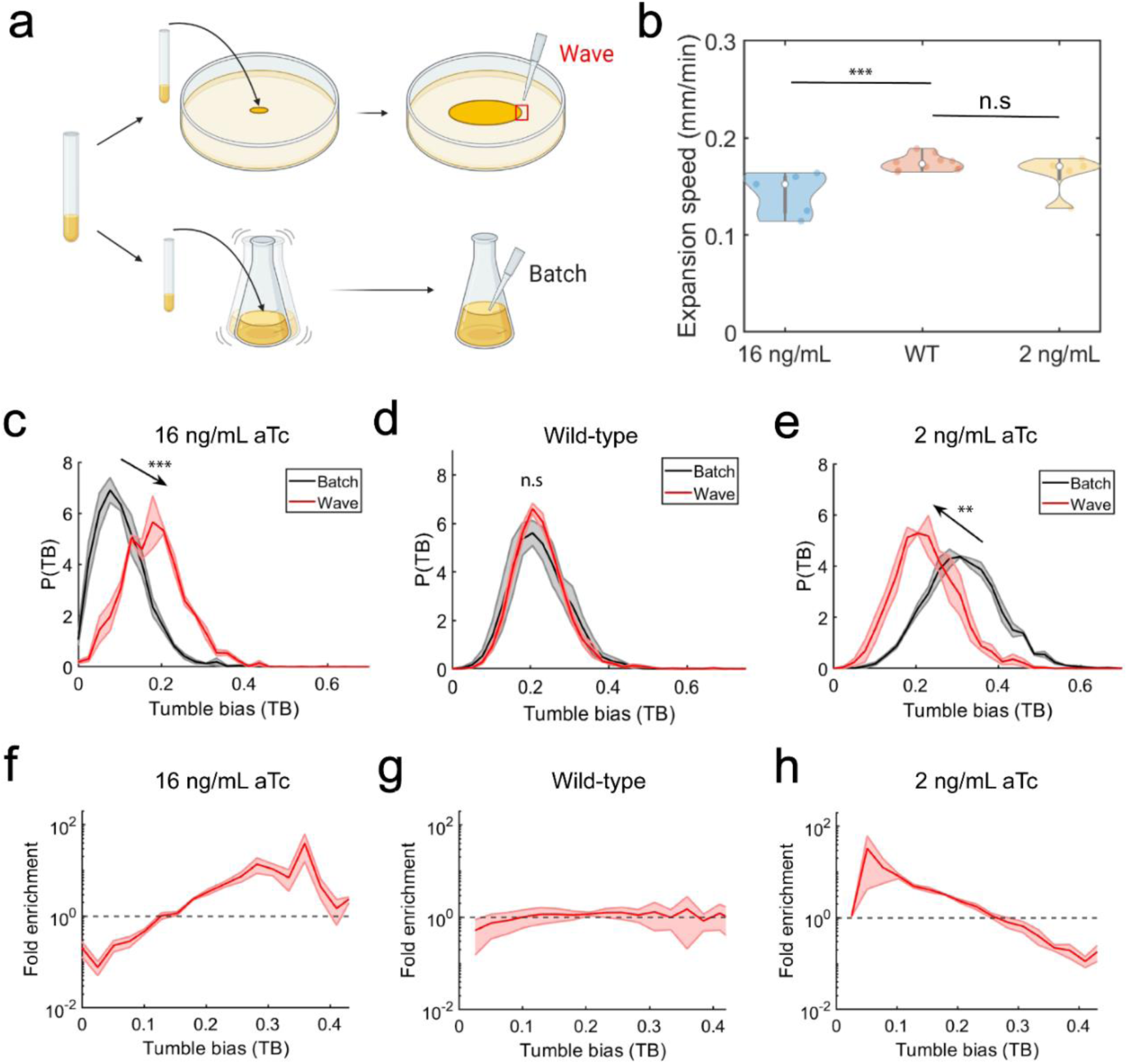
Migration in agarose swim plate tunes TB distribution of the population. **(a)** Experimental design: a swim plate and shaking flask containing the same amount of inducer are inoculated with the same initial population. TB distributions are measured and compared between batch and the migrating ring (“wave”) after spending the same amount of time in either condition. **(b)** Expansion speeds of RP437 wild-type (mean expansion speed 0.174 ± 0.007 mm/min; uncertainties throughout the paper are the standard errors of the mean across replicates; here, *N*_*rep*_ = 12, ΔcheZ (pTet-CheZ) induced with 2 ng/mL aTc (mean 0.170 ± 0.02 mm/min; *N*_*rep*_ = 6), and 16 ng/mL (mean 0.150 ± 0.02 mm/min; *N*_*rep*_ = 6) populations on agarose swim plates. The expansion radius over time was measured laterally and in one direction from the center to the edge of the migrating colony. Two-sided t-tests were performed throughout the text to determine significance of the difference between pairs of means (throughout, *** = P<0.0005, ** = P<0.005, * = P<0.05, and n.s. = not significant). **(c)** Populations with low initial mean TB (mean TB = 0.10 ± 0.01; *N*_*rep*_ = 3, *N*_*cells*_ = 21,891) shift toward a higher mean TB after migration on the plates (0.18 ± 0.01; *N*_*rep*_ = 3, *N*_*cells*_ = 5,103). Shaded area on the TB distribution is the standard error of the mean probability in each TB bin. Significance here and throughout the figures refers to change in the mean of the distribution. **(d)** There was minimal or no shift in wild-type population’s TB distribution during collective migration on the plates (0.22 ± 0.01; *N*_*rep*_ = 3, *N*_*cells*_ = 15,774 *cells*) compared to batch culture (0.22 ± 0.03; *N*_*rep*_ = 3, *N*_*cells*_ = 9,417). **(e)** On the other hand, populations with high initial mean TB (0.31 ± 0.01; *N*_*rep*_ = 3, *N*_*cells*_ = 26,254) shifted toward a lower mean TB after migration (0.23 ± 0.03; *N*_*rep*_ = 3, *N*_*cells*_ = 6,951). **(f)** When low-TB populations migrated on the plates, fold enrichment of cells with high TB is observed, while cells with low TB are filtered out (*N*_*rep*_ = 3). Shaded area on the fold-enrichment throughout is the standard errors of the mean enrichment values within each TB bin **(g)**. No enrichment and filtering of TB were observed in wild-type populations (*N*_*rep*_ = 3). **(h)** When high-TB populations migrated on the plates, enrichment of cells with low TB is observed, while cells with high TB are filtered out (*N*_*rep*_ = 3).

The inducer aTc degrades over time; therefore, we aged aTc before adding it to the batch culture growth medium to mimic the conditions in the migrating wave (**Fig. S2; Methods**). The same experiments were performed with WT cells in the absence of aTc. By comparing the TB distributions of populations that had migrated on the plates with those propagated in batch culture, we quantified the effect of collective migration on the populations’ TB distributions.

We found that the TB distribution of the high-TB population (batch mean TB = 0.31 ± 0.01, uncertainties throughout are the standard errors of the mean across biological replicates) shifted toward lower TBs (final mean TB = 0.23 ± 0.03) during migration on plates (**Fig. 2e, h**). Conversely, the TB distribution of the low-TB population (batch mean TB = 0.10 ± 0.01) shifted to higher TBs (final mean TB = 0.18 ± 0.01) after migration on the plates (**Fig. 2c, f**). In both cases, the populations became enriched for cells with intermediate TBs of ∼0.2, close to the mean TB of wild-type populations grown in batch culture (mean TB = 0.22 ± 0.03) (**Fig. S3**). Consistent with these observations, the TB distribution of the migrating WT population (final mean TB = 0.22 ± 0.01) was similar to its batch culture distribution (**Fig. 2d, g**). All changes in the mean TB during migration were statistically significant, as measured by two-sided t-test, except that of the WT population. Samples harvested 1 cm behind the migrating populations were marginally enriched for the low-performing TBs, though the effect was not significant, possibly because growth also occurred between the moment when these cells fell behind and when they were harvested (**Fig. S4**).

To quantify how these shifts in phenotype composition affected migration speed, we measured the expansion radius of the migrating waves of the three populations on agarose swim plates over time and calculated the steady-state expansion speed using the slope of the linear part of the expansion radius profile (**Methods;** Fig. 2b and **Fig. S5**). Early time points in migration could not be measured because the ring was not clearly visible. We found that the CheZ mutants reached their steady state speeds much later than the WT, possibly due to the scarcity of high-performing phenotypes in the batch culture distributions. However, once the CheZ mutants reached steady state, their migration speeds were close to that of the WT, illustrating the effectiveness of the adaptation mechanism. In particular, the WT population (mean steady-state expansion speed 0.174 ± 0.007 mm/min) and the high-TB population (0.170 ± 0.02 mm/min) expanded the fastest (difference in speed was not statistically significant). The low-TB population was slower (0.150 ± 0.02 mm/min; significant to P < 0.005), possibly due to the presence of very low-TB cells in the adapted population (**Fig. S3**). The growth rates of WT, low-TB, and high-TB populations were similar (**Fig. S6**), therefore differences in the steady state expansion speeds were solely due to differences in chemotactic performance. Thus, migrating populations become enriched with high-performing phenotypes, increasing migration speed. In this environment, cells with TB close to 0.22 appear to perform best, which coincides with the average TB of the WT population in batch culture (**Fig. S3**).

### Tuning of swimming phenotype distributions is non-genetic and occurs without changes in *cheYZ* expression

What mechanism enables a migrating bacterial population to tune its TB distribution? Our recent theory predicts that the tuning of phenotypic distributions is a dynamic, non-genetic process mediated by the loss of low-performing chemotaxis phenotypes^15^ and production of new phenotypes through growth^18,24,25^. However, during our swim plate assays, cells in the migrating populations could potentially acquire mutations that permanently altered their swimming behaviors^4–6^. If TB tuning was non-genetic, then we expected that the TB distributions of cells in the migrating waves would revert to their standing TB distributions when grown in batch culture.

To distinguish between these possibilities, we isolated cells from the edge of the low- and high-TB migrating colonies on agarose swim plates, inoculated them into batch cultures with aTc, and measured their TB distributions after growth (**Fig. 3a**).

**Figure 3:**
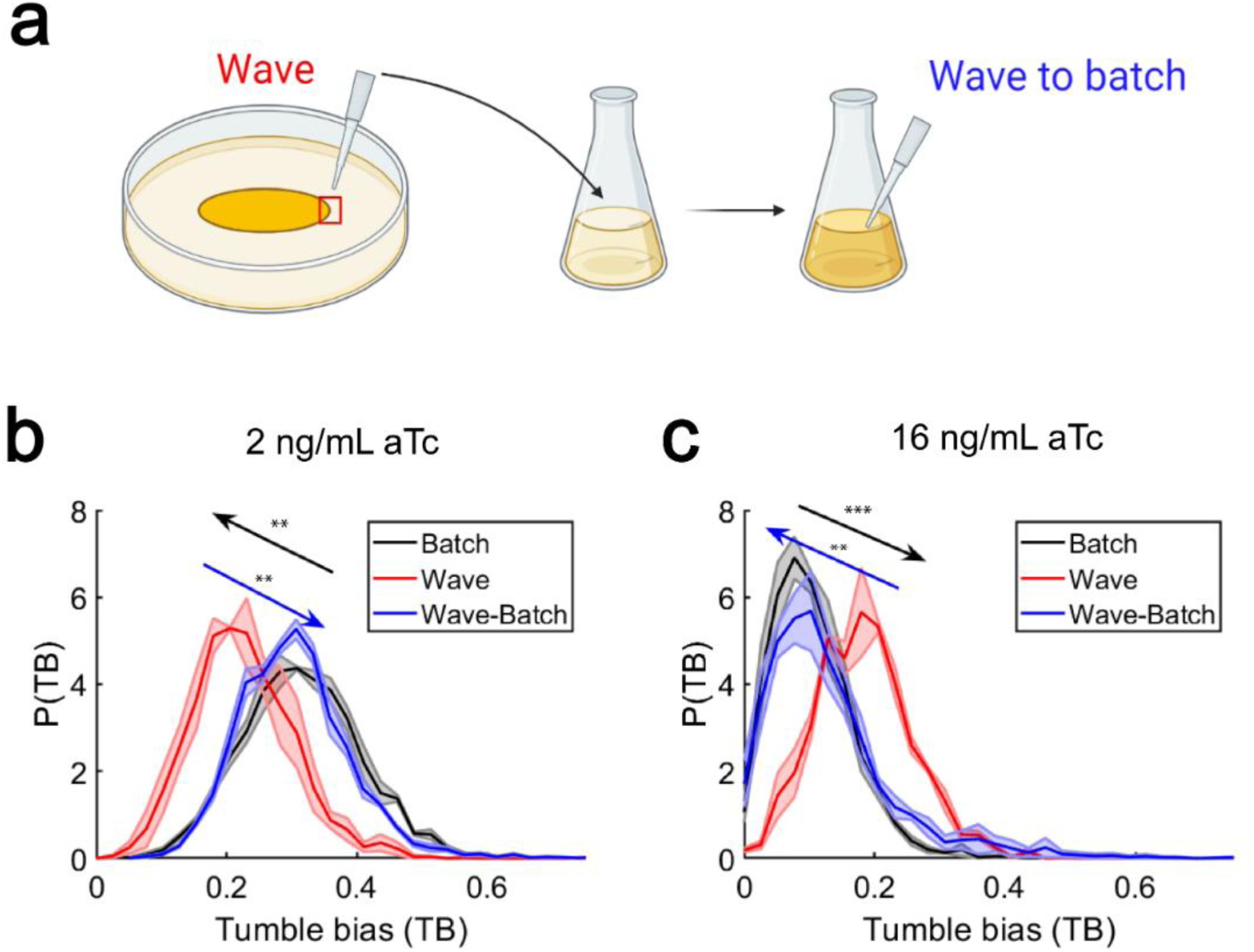
Tuning of TB by collective migration is reversible. **(a)** To determine whether the tuning of TB becomes fixed or remains adaptable, we isolated cells from the edge of the migrating colonies (“wave”), grew the cells in liquid batch culture until exponential phase, and measured the TB distributions after growth (“wave-batch”). **(b)** Black and red are the TB distributions of populations with high initial TB before and after migrating on agarose swim plates (data replotted from **Fig. 2e**). Blue: TB distribution of cells that migrated (red) after regrowth in batch culture (mean TB = 0.30 ± 0.01; *N*_*rep*_ = 3, *N*_*cells*_ = 15,774). The TB distributions of the populations that migrated on agarose swim plates, after growing in flasks, shifted back to the original batch culture’s distributions. **(c)** Similar to B, the TB distributions of populations with low initial TB, after growing in flask (blue; 0.13 ± 0.02; *N*_*rep*_ = 3, *N*_*cells*_ = 20,403), shifted back to the TB distributions observed in batch culture (black and red distributions are replotted from **Fig. 2c**). The mean values of “batch” and “wave-batch” TB distributions in both cases are not significantly different.

The TB distributions of the migrating populations relaxed to their standing TB distributions after re-growth in batch culture, suggesting that the cells in the migrating population did not acquire mutations that permanently altered their swimming behaviors (**Fig. 3b, c**).

Even without mutations, the observed shifts in TB distribution during migration could have been due to changes in gene expression. Chemotaxis is regulated during *E. coli* growth, leading to changes in a population’s TB distribution at different growth phases^24^. If cells experienced different growth conditions as they migrated, expression of the response regulator CheY and its phosphatase CheZ could be affected, thus changing TB^15,17^. Given that there was excess nutrient on the plates, we expected that bacteria would remain in exponential phase for the duration of the experiment, and therefore that *cheY* and *cheZ* regulation would be constant throughout the plate as the population migrates. We measured the protein expression levels of CheY and CheZ by fusing them to fluorescent proteins (RFP and YFP, respectively) at their native chromosomal loci in a non-motile strain. This non-motile strain acted as a biosensor for detecting *in vivo* changes in *cheYZ* expression. We spread these cells uniformly on agarose swim plates, initiated migrating waves of motile, unlabeled WT cells at the center of the plates, and then picked cells from both the center and the edge of the migrating colonies (**Methods**). Fluorescent microscopy measurements (**Methods**) revealed no significant differences in CheY and CheZ levels between cells sampled from the center vs. the edge (**Fig. S7**). This indicated that the expression of *cheY* and *cheZ* remains constant across the entire plate even as the wave migrates. We further confirmed that migrating populations did not experience catabolite repression^26,27^ during the period of our measurements since catabolite repression effects only appeared at much longer times (**Fig. S8**).

Together, these observations indicated that the adaptation of TB distributions was highly unlikely to be mediated by mutations or gene regulation.

### Tuning of swimming phenotype distributions adapts the population to the physical environment

We have demonstrated that migrating cell populations non-genetically tune their own phenotypic composition, enriching the population for high-performing chemotaxis phenotypes. Since the performance of chemotaxis phenotypes varies in different environments^18^, we expected that the TB distribution during migration would depend on the environment that the population traversed. Whereas cells need to tumble to escape traps in semisolid agarose^18,23,28^, past work has shown that cells with low TB climb chemical gradients faster in liquid^17,29^. This suggests that the highest-performing phenotype in liquid has a lower TB than that in semisolid agarose, and thus migrating populations would enrich for lower TBs in liquid than in agarose.

To test this, we measured the TB distributions of migrating populations after traveling through a liquid environment (**Fig. 4a**). We again generated populations of *E. coli* with low and high average TBs and propagated migrating waves with these populations, as well as the wild-type, in capillary tubes filled with liquid media for 9 hours. aTc was pre-aged so that the total aging time at the end of the capillary experiments was the same as in our agarose swim plate experiments. We then isolated cells from the migrating populations and measured their TB distributions after migration.

**Figure 4.**
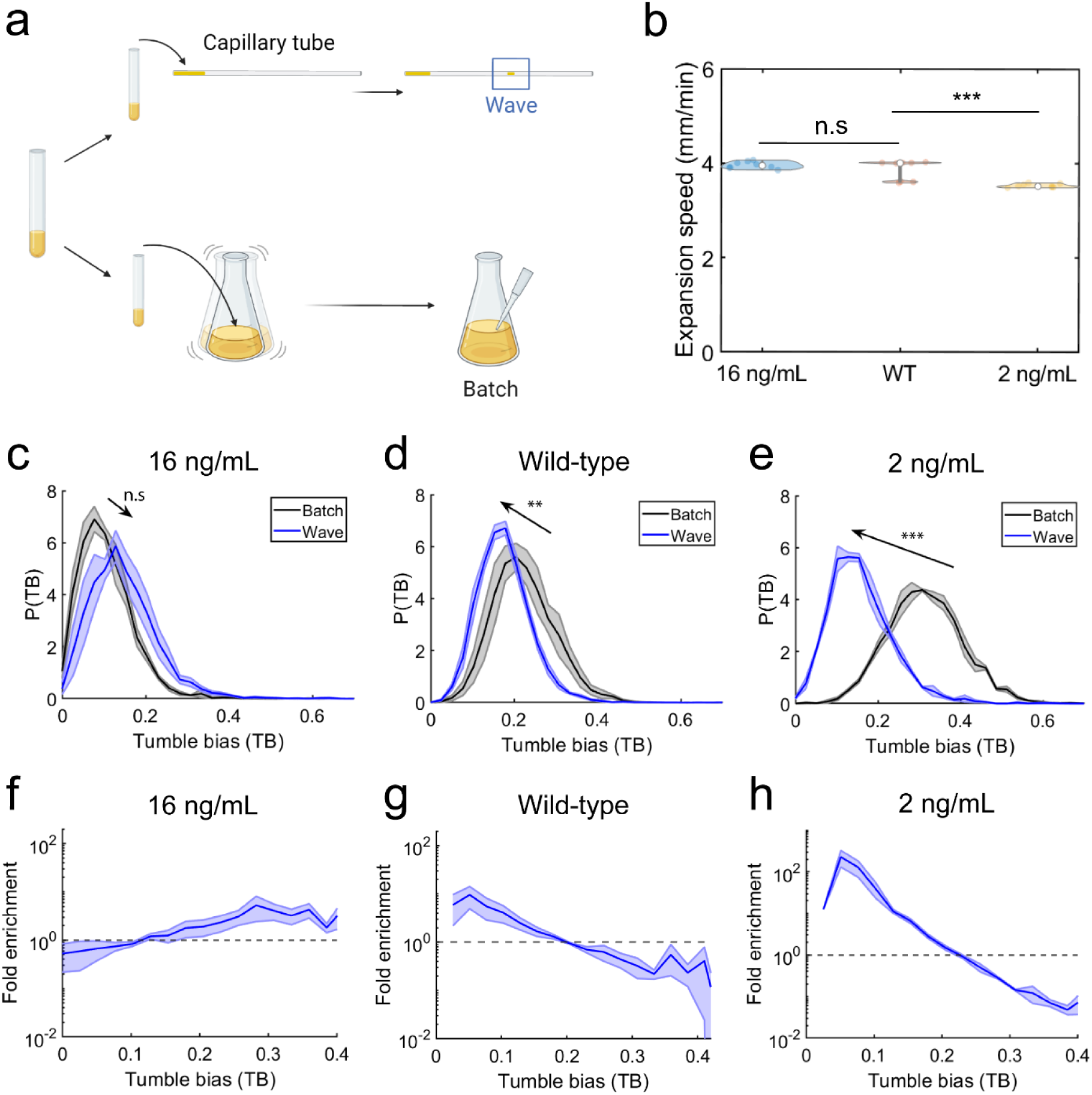
Tuning of TB distribution is different in liquid environment: **(a)** Experimental design: a capillary tube and shaking flask containing the same amount of inducer are inoculated with the same initial population. TB distributions are measured and compared between batch and the migrating cells (“wave”). **(b)** Expansion speeds of RP437 wild-type (mean 3.86 ± 0.02 mm/min; *N*_*rep*_ = 9, ΔcheZ (pTet-CheZ) induced with 2 ng/mL aTc (mean 3.53 ± 0.04 mm/min; *N*_*rep*_ = 9), and 16 ng/mL aTc (3.96 ± 0.01 mm/min; *N*_*rep*_ = 9) populations in capillary tubes. **(c)** In liquid, populations with low initial mean TB (mean TB = 0.10 ± 0.01; *N*_*rep*_ = 3, *N*_*cells*_ = 21,891) shift toward a higher mean TB (0.15 ± 0.03; *N*_*rep*_ = 3, *N*_*cells*_ = 20,403), but which is slightly lower than the mean TB in agarose swim plates (0.18 ± 0.01, as in **Fig. 2c**). **(d)** Unlike in agarose swim plates, wild-type population’s TB distribution (0.22 ± 0.03; *N*_*rep*_ = 3, *N*_*cells*_ = 9,417) shifted toward a lower mean TB during collective migration in capillary tubes (0.18 ± 0.01; *N*_*rep*_ = 3, *N*_*cells*_ = 16,768). **(e)** Populations with high initial mean TB (0.31 ± 0.01; *N*_*rep*_ = 3, *N*_*cells*_ = 26,254) shifted toward a lower mean TB (0.16 ± 0.01; *N*_*rep*_ = 3, *N*_*cells*_ = 13,434). The mean TB value is lower than what was observed during migration on agarose swim plates (0.23 ± 0.03, as in **Fig. 2d**). **(f)** For the migration of low-TB populations, the fold enrichment is smaller than in agarose swim plates. (*N*_*rep*_ = 3) **(g).** Wild-type populations migrating in capillary tubes were enriched for low-TB cells (*N*_*rep*_ = 3), unlike what we observed when they migrated on agarose. **(h)** For high-TB populations, fold enrichment was stronger in capillary tubes compared to migration in agarose swim plates (*N*_*rep*_ = 3). More low-TB cells are enriched during migration in capillary tubes than on agarose plates.

The WT population’s TB distribution (batch mean TB = 0.22 ± 0.03) shifted to lower TB (final mean TB = 0.18 ± 0.01) during collective migration in liquid (**Fig. 4d, g**). Low-TB cells were enriched up to a factor of 10, whereas high-TB cells were filtered out. When the population with high TB (batch mean TB = 0.31 ± 0.01) traveled through capillary tubes, its TB distribution shifted lower than it did when traveling on agarose swim plates (final mean TB = 0.16 ± 0.01) (**Fig. 4e**). The enrichment of low-TB cells and filtering out of high-TB cells were much stronger in the capillary tubes (**Fig. 4h**) compared to the agarose swim plates (**Fig. 2h**). The TB distributions of the low-TB population (batch mean TB = 0.10 ± 0.01) shifted to a slightly higher mean TB when traveling through capillary tubes (final mean TB = 0.15 ± 0.03), but the shift was not statistically significant (**Fig. 4c**). In this case, the enrichment of high-TB cells and filtering out of low-TB cells was weaker in the capillary tubes (**Fig. 4f**) compared to agarose swim plates (**Fig. 2f**).

As in agarose, adaptation enabled the CheZ mutants and WT population to reach similar steady state migration speeds. The low-TB population had the fastest expansion speed (mean expansion speed 3.96 ± 0.01 mm/min). Its TB distribution shifted the least during migration in capillary tubes, suggesting that its standing TB distribution was better adapted for migration in liquid than the WT and high-TB populations (**Methods; Fig. 4b**). The expansion speed of the WT population was statistically indistinguishable from this (3.86 ± 0.02 mm/min). Finally, the high-TB population expanded slower than the other two populations (3.53 ± 0.04 mm/min).

Together, these results established that shifts in TB distribution during collective migration of *E. coli* enriched for phenotypes with highest chemotaxis performance in the environment being traversed.

We also tested whether environment-dependent adaptation of TB distributions by collective migration occurred in other wild-type *E. coli* strains, including: 1) HE205, a NCM3722 derivative from the Cremer and Hwa labs^21^, 2) AW405, a K-12 strain originated from the Berg lab^23^, 3) MG1655, a common K-12 lab strain^30^, and 4) TW09231, or *Escherichia sp.* TW09231, a wild isolate from Lake Michigan^31^. In all cases, migration in liquid resulted in populations enriched with lower TB cells than when migrating in agarose (**Fig. S9**), consistent with adaptation by collective migration. These results demonstrate environment-dependent tuning of TB distributions across a wide range of both lab *E. coli* strains and a wild isolate, highlighting the universality of this adaptation mechanism.

### Adapted TB distribution rapidly relaxes back to batch distribution

How quickly does the adapted phenotypic distributions of a migrating population reverse back to the standing batch distribution? To quantify this, *E. coli* populations were isolated from capillary tube assays and reinoculated into batch cultures. TB distributions were measured every 15-30 minutes throughout the growth period (**Fig. 5a**). We fitted the mean TB values over time to an exponential model to determine the characteristic timescale on which the mean TB returns to the level observed in batch culture (**Methods**). For wild-type RP437, it took about 2 doubling times for the TB distributions to return to those observed in batch cultures (**Fig. 5b**). The relaxation time scale of the mean was *τ* = 65.2 ± 1.6 min and the doubling time was 68.8 ± 1.5 min. The result was similar for an *E. coli* strain with slower growth rate, HE205. Its TB distribution also relaxed within approximately 2 doubling times (**Fig. 5c**). The relaxation time scale of the mean was *τ* = 106.1 ± 6.3 min and the doubling time was 92.3 ± 3.4 min. These results demonstrate that adaptation by collective migration is rapidly reversible, requiring only a few generations of growth.

**Figure 5.**
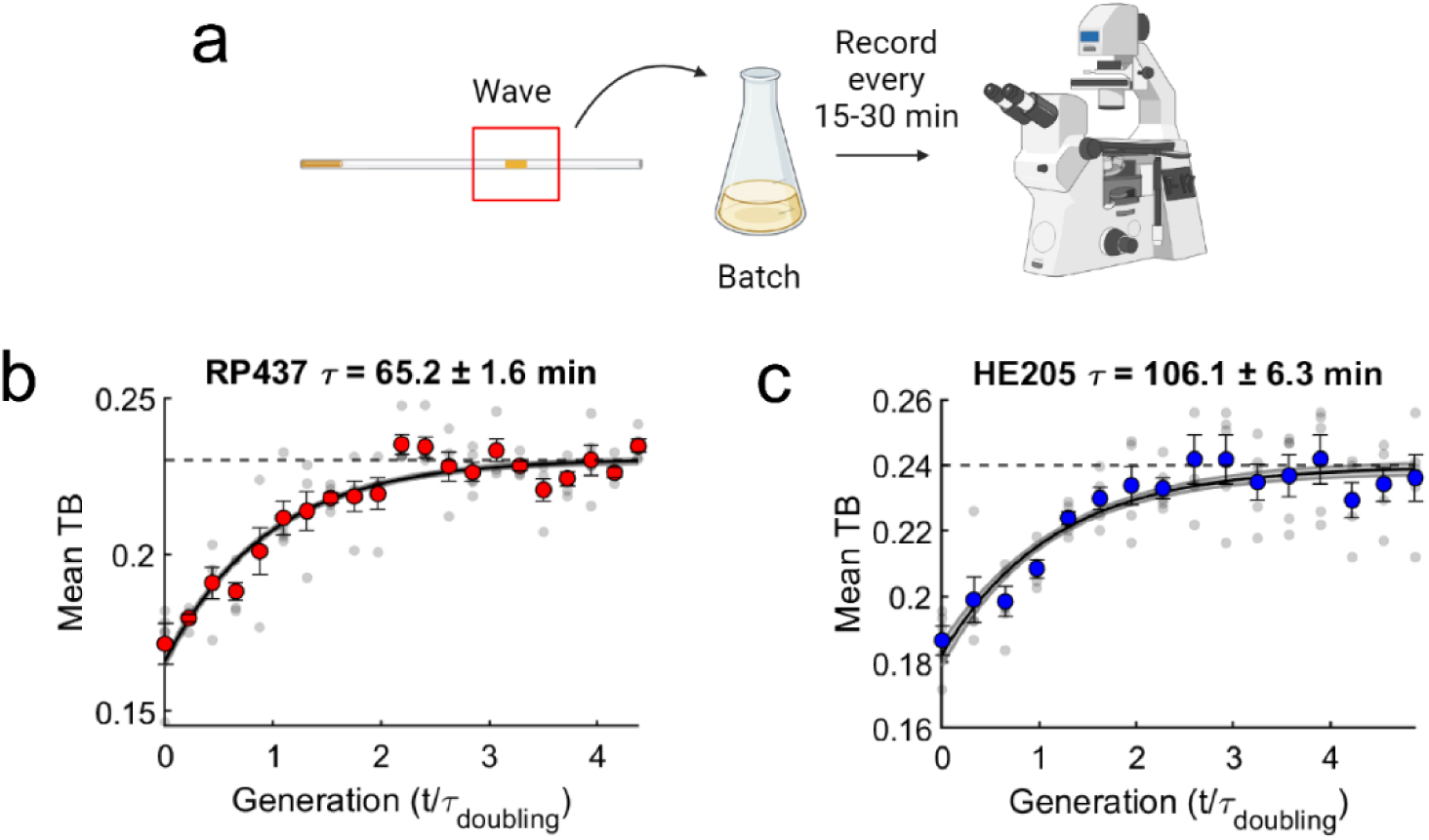
Adapted TB distribution relaxes back to the standing batch distributions within a few generations. (**a**) Experimental design: E. coli RP437 and HE205 populations were inoculated in capillary assays and let to migrate for 9 hours. Migrating populations were isolated and grown in batch cultures. TB measurements were taken from the batch culture over the course of growth every 15 minutes (RP437) or −1 30 minutes (HE205). (**b**) In ∼2 generations (growth rate = 0.604 ± 0.01 hour^−1^; *τ*_*doubling*_ = 68.8 ± 1.5 minutes; errors are the standard errors of the mean growth rates and doubling times; *N*_*rep*_ = 3), the mean TB of migrating populations returns back to the mean TB of populations observed in batch culture (the mean TB of the batch culture is represented as a horizontal dashed line). The gray dots are the mean TB of individual biological replicates. The red dots are the means across replicates (calculated from more than 30,844 trajectories for each time points; *N*_*rep*_ = 5), and the error bars are the standard errors of the mean. The means and standard errors across replicates were fitted with an exponential model, and a characteristic relaxation timescale *τ* and its error were determined using maximum likelihood estimation (see **Methods**). The black line is the model fit, and the shaded area is the standard deviation of the model. *τ* for RP437 is 62.5 ± 1.6 minutes – the error is the standard deviation of the estimated *τ*.(**c**) Same as **b**, but for HE205. The blue dots are the means across replicates (calculated from more than 5,920 trajectories each time −1 points; *N*_*rep*_ = 5). It takes ∼2 generations (growth rate = 0.45 ± 0.01 hour; *τ*_*doubling*_ = 92.3 ± 3.4 minutes; *N*_*rep*_ = 3) for the TB distributions to relax back to the batch culture. *τ* for HE205 is 106.1 ± 6.3 minutes.

### Collective migration tunes the population’s distribution of sensing phenotypes to adapt to the chemical environment

Numerous phenotypic traits, in addition to tumble bias, affect a bacterium’s chemotactic performance. One such trait is the cell’s sensing capability, which correlates with the abundances of its chemoreceptors^32,33^. *E. coli* express five chemoreceptor species, of which ∼90% are either Tar, which binds preferentially to L-aspartate, or Tsr, which binds preferentially to L-serine^34–36^. Notably, Tsr is expressed separately from the chemotaxis operon—which includes, among other genes, Tar, CheY, and CheZ^37^—allowing its abundance to vary independently of other chemotaxis genes affecting chemotaxis performance. Therefore, we predicted that Tsr could be enriched in a migrating wave that chases its cognate ligand L-serine.

We measured the Tsr protein expression levels in populations of *E. coli* migrating on agarose swim plates with 100 µM L-aspartate or 100 µM L-serine as attractants, and no other amino acids (**Fig. 6a**). As a carbon source, we used glycerol because it is not an attractant^38^. To measure single-cell Tsr levels, we fused *tsr* on its native locus with mYFP (**Methods**). We used MG1655 as the parent strain here because, unlike RP437^37,39^, it is not restricted by auxotrophic limitation and can grow in minimal media in the absence of amino acids. The expansion speed of the population was faster in serine (0.88 ± 0.05 mm/hr) than in aspartate (0.52 ± 0.1 mm/hr) (**Fig. 6b**) consistent with previous reports^21^.

**Figure 6:**
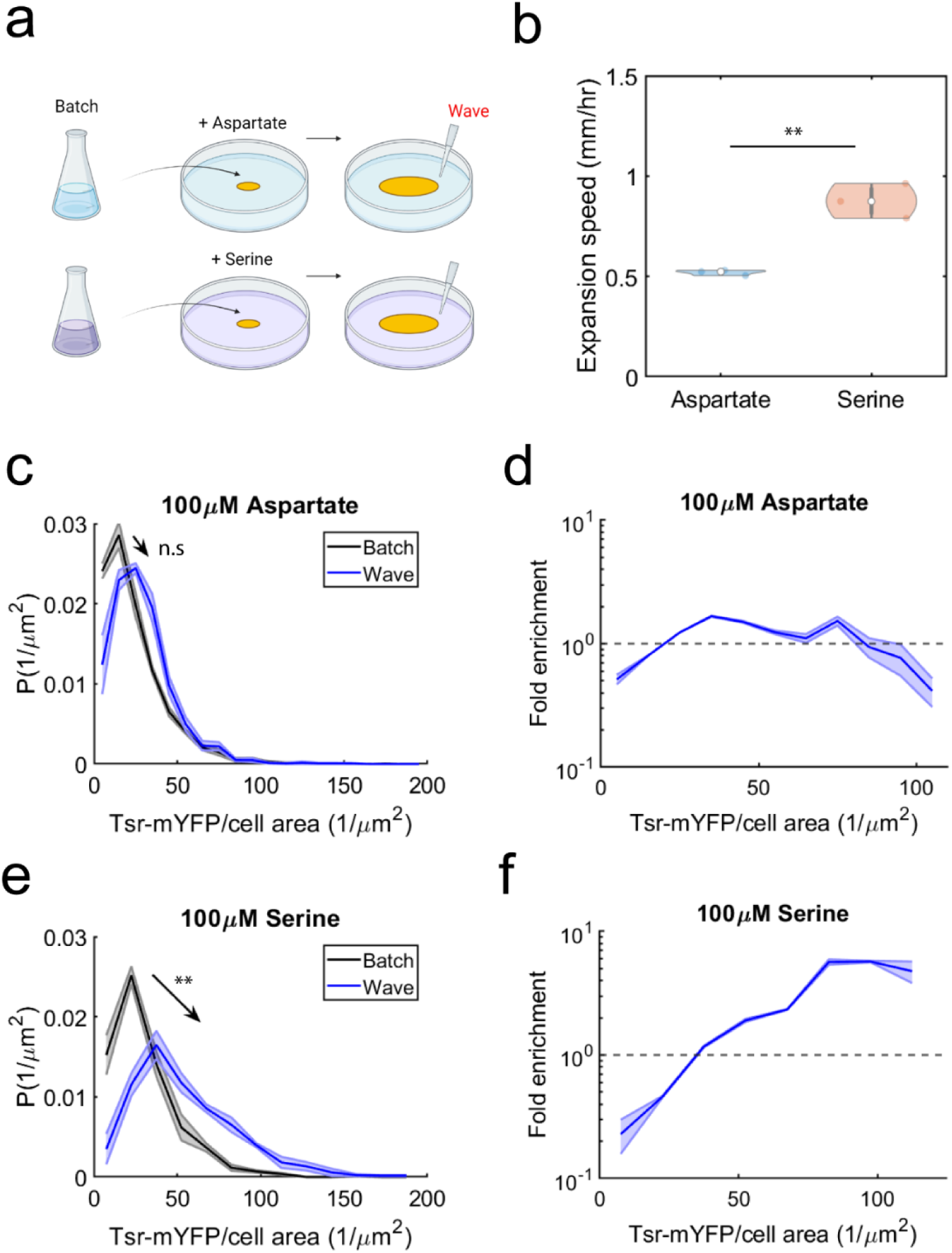
Collective migration tunes the chemoreceptor levels depending on the attractants. **(a)** Experimental design: E. coli MG1655 wild-type with YFP-labelled Tsr was inoculated on swim plates supplemented with 100 μM aspartate or 100 μM serine. The fluorescence of single cells was measured in populations at the edge of the migrating colonies (“wave”) and in populations growing in batch culture. The single-cell fluorescence divided by the cell area is used as a proxy for receptor concentration. (**b**) The expansion speeds of the migrating populations in agarose swim plates supplemented with 100 μM aspartate (0.52 ± 0.1 mm/hr; *N*_*rep*_ = 3) or 100 μM serine (0.88 ± 0.05 mm/hr; *N*_*rep*_ = 3). (**c**) Black is the distribution of Tsr levels of populations grown in batch culture with added aspartate (mean Tsr intensity = 23.7 ± 0.4 *µm*^−2^; *N*_*rep*_ = 3, *N*_*cells*_ = 2,076), and blue is the distribution of Tsr levels of the migrating populations that chased aspartate (27.2 ± 3.8 *µm*^−2^; *N*_*rep*_ = 3, *N*_*cells*_ = 1,617). Shaded areas on the Tsr intensity distributions are the standard errors of the mean probabilities in each bin of intensity value. (**d**) When chasing aspartate, the distributions of Tsr levels in migrating populations remained close to that in batch cultures, with only a weak fold increase in cells with intermediate Tsr levels and a weak fold decrease in cells with high Tsr levels. Shaded area on the fold-enrichment throughout is the standard errors of the mean enrichment values within each bin of Tsr intensity value. (**e**) Black is the distribution of Tsr levels of populations grown in batch culture with added serine (29.6 ± 2.9 *µm*^−2^; *N*_*rep*_ = 3, *N*_*cells*_ = 1,695), and blue is the distribution of Tsr levels of the migrating populations that chased serine (53.6 ± 6.5 *µm*^−2^; *N*_*rep*_ = 3, *N*_*cells*_ = 675). (**f**) During collective migration on swim plates supplemented with serine, there was a strong enrichment of cells with higher Tsr levels, while cells with lower Tsr levels were filtered out.

We compared the average mYFP fluorescence in single cells picked from the edge of the migrating colonies versus those grown in batch cultures supplemented with the same concentration of either L-aspartate or L-serine (**Methods**). When aspartate was the chemoattractant, distributions of Tsr levels within the migrating populations on swim plates were similar to those grown in batch cultures, with the mean Tsr level statistically indistinguishable from batch culture (**Fig. 6c**; mean intensities 〈Tsr〉_batch_ = 23.7 ± 0.4 μm^−2^ and 〈Tsr〉_wave_ = 27.2 ± 3.8 μm^−2^). We observed a slight enrichment of intermediate Tsr levels (**Fig. 6d**), possibly because of the assistance Tsr provides to the adaptation of Tar (CheR and CheB can methylate and demethylate neighboring Tar receptors when bound to the pentapeptide of Tsr)^40,41^. In contrast, collective migration led to strong enrichment for cells with high levels of Tsr when the population chased serine (**Fig. 6e, f**; 〈Tsr〉_batch_ = 29.6 ± 2.9 μm^−2^ and 〈Tsr〉_wave_ = 53.6 ± 6.5 μm^−2^). While the distribution of Tsr changed significantly depending on the attractant the population was chasing (see **Fig. S10** for a direct comparison), the distributions of TB were similar across attractants, with only a mild adjustment in serine (**Fig. S11**), suggesting that adaptation of these two traits is nearly independent.

To confirm that the shift in the distribution of Tsr levels was not due to gene regulation, we employed an analogous strategy to the one described above for CheY and CheZ (**Fig. S7a**), here using a non-motile strain expressing fluorescently-labelled Tsr from its native locus (**Fig. S12a**). There were no differences in the Tsr levels between non-motile cells harvested at the migrating edge and at the center of the plate, suggesting that *tsr* gene expression changes negligibly during collective migration (**Fig. S12b, c**). Tsr levels were not regulated by serine or aspartate consumption (when these amino acids were present in batch cultures at the same concentrations as in the swim plates) (**Fig. S13**), nor were TB distributions (**Fig. S14**). This further indicated that the receptor enrichment that we measured after migration on swim plates was not caused by altered gene expression or metabolic effects due to the presence of amino acids.

Taken together, these observations demonstrated that adaptation by collective migration generalizes to sensory phenotypes: migrating populations non-genetically adapted their distributions of receptor abundances, depending on the attractants they were chasing. Thus, collective migration combined with growth can enrich for multiple, continuous phenotypic traits and non-genetically adapt migrating populations to diverse environmental conditions.

## Discussion

Here, we investigated how a population of cells adapts its distribution of phenotypes to migrate effectively in multiple environments. We discovered that migrating populations of *E. coli* can tune their distributions of swimming behaviors and sensing capabilities based on the physical and chemical properties of their environment. This tuning occurs dynamically and is explained by a model in which loss of low-performing chemotaxis phenotypes from a migrating front^15^ balances the production of new phenotypes by growth^18,25^. This mechanism, which we demonstrated in multiple *E. coli* strains, can act on pre-existing variability in gene expression but does not require *de novo* changes in gene expression or mutations. By rapidly and reversibly enriching for fitter individuals based on the demands of the current environment, this mechanism increases the population’s expansion speed across a wide variety of potential environments.

We examined two phenotypes that impact chemotaxis performance: tumble bias and receptor abundance. In liquid, low tumble bias cells are known to climb a fixed gradient faster^17^. Consistent with this wild-type RP437 populations enriched for low-TB cells when migrating in liquid. When migrating on agarose, however, the TB distribution did not change from the standing batch distribution. Furthermore, both CheZ-inducible populations, with high or low initial mean TB, shifted toward the standing batch distribution of the wild-type RP437 (**Fig. S2**). This suggests that the standing batch TB distribution of wild-type RP437 is already adapted for navigation in agar/agarose. Indeed, RP437 is a standard laboratory strain that has been used to study motility for decades^39,42^ and was originally selected for motility on agar swim plates. In all the strains we tested, the tumble bias distribution when migrating in liquid was enriched for lower TB cells than when migrating on agarose (**Fig. 5a, b**, and **Fig. S9**). This is consistent with previous findings that tumbling slows migration in liquid^17,29^, but that tumbles are needed to escape traps in porous environments ^23,43,44^. More broadly, we expect that any phenotypic trait that contributes to chemotaxis performance can be enriched during collective migration. Furthermore, it is always possible for one of the countless strains of bacteria in nature to encounter an environment in which its phenotype composition is poorly adapted. In these cases, we expect non-genetic adaptation to have large effects on its traveling phenotype distribution.

We also found that the adapted TB distributions relaxed back to the batch culture distribution in about two generations, indicating some degree of nongenetic inheritance. This is consistent with a previous study which showed that swimming traits of individual *E. coli* cells are partially and nongenetically inherited across multiple generations^25^. While non-genetic inheritance of chemotactic traits^25^, and other traits like antibiotic resistance^45^, sporulation^46^, iron metabolism^47^ were studied previously, the molecular mechanism of how these traits are inherited remains to be explored further. Indeed, non-genetic inheritance of chemotactic traits may involve partitioning of chemoreceptors, flagellar components, and other regulatory elements from the mother to daughter cells during cell division.

Recent research emphasizes the crucial role of emergent spatial structures during collective migration in enhancing genetic diversity within bacterial communities^48^. In traditional well-mixed environments, resource competition leads to the extinction of species with lower relative growth fitness^49^. However, during collective migration, species with lower relative growth fitness can coexist by competing for spatial territories rather than resources^4,48^, and avoid extinction by colonizing specific habitats within the migrating colony. Furthermore, recent experiments demonstrate the evolution of mutants with different navigation capabilities from isogenic migrating populations^4^, suggesting that the emergent spatial organization of phenotypes could contribute to genetic diversity^15,21,50,51^. Together with our observation that environment determines the specific distribution of phenotypes that is enriched, these results suggest that relative spatial arrangements of phenotypes within migrating groups may play an important role in eco-evolutionary dynamics.

Collective migration is not exclusive to bacteria; it also occurs in eukaryotic systems where cells consume or degrade attractants to form and chase self-generated gradients, a mechanism observed in various contexts including cancer and development^52,53^. Some examples are the migration of lateral line of zebrafish embryos^54^, neural crest cells in *Xenopus laevis*^55^, and melanoma cells^56^. The tuning of phenotypic distributions during collective migration only requires that some cells, due to their specific traits, are more adept at navigating than others, therefore creating a “leader-follower” structure. Thus, adaptation by collective migration may be a general strategy for cell populations to colonize diverse environments effectively.

Our work demonstrates a new mechanism for population-level adaptation in biology. Adaptation mechanisms such as stochastic switching^7,9,57^ and regulating gene expression are fast (∼1 generation) but are limited to the adaptation of a few phenotypic traits and often require dedicated pathways to implement. Adaptation by genetic mutations avoids these limitations but is slow (∼10 generations)^4–6^. In contrast, standing variation in numerous phenotypic traits among individual cells is ubiquitous. Our findings demonstrate that when collective behaviors create selection pressures which shape that variation, cell populations can reversibly adapt multiple traits with a level of speed and flexibility that is difficult to achieve via classical mechanisms.

## Methods

### Strains, media, and growth conditions

Most experiments were performed with *E. coli* K12 RP437 and MG1655 derivatives. Other strains used in this study are *E. coli* HE205, AW405, and wild-isolate TW09231 (*Escherichia sp.* TW09231). To tune tumble bias distributions, we utilized *E. coli* RP437 *ΔcheZ* and HE205 *ΔcheZ* strains that carry an aTc-inducible cassette of *cheZ* that is integrated in the chromosome at the *attB* site (RP437 *ΔcheZ pTet-cheZ* and HE205 *ΔcheZ pTet-cheZ*). To measure single cell Tsr receptor abundances, we utilized an *E. coli* MG1655 *ΔfliC:FLP* mutant (gift from Victor Sourjik) that has *tsr* translationally fused with monomeric YFP (A206K) on its native chromosomal locus (MG1655 *ΔfliC:FLP, tsr*-*mYFP*). To modulate motility in the previous strain, we transformed an arabinose-inducible plasmid expressing WT FliC (*pBad*-*fliC*, gift from Howard Berg) into MG1655 *ΔfliC:FLP, tsr-mYFP* and induced it with 0.005% w/v arabinose. RP437 *ΔfliC* with CheY and CheZ translationally fused with mRFP and mYFP, respectively on their native chromosomal loci (RP437 *ΔfliC cheY-mRFP and cheZ-mYFP*) was used as a biosensor to measure expression of the *cheY* and *cheZ* genes in the swim plate assays. MG1655 *ΔfliC:FLP*, *tar*-*mCherry, tsr*-*mYFP*, was used as a biosensor to measure expression of the *tsr* gene. To insert genes on the native loci of the *E. Coli* chromosome, we employed the phage λ Red recombination system.

RP437-derived strains were stored as −80°C freezer DMSO-based stocks, and MG1655-derived strains were stored as glycerol-based stocks at the same temperature. M9 buffer supplemented with casamino acids, glycerol, and MgSO_4_ (1x M9 salts, 0.4% v/v glycerol, 0.1% v/v casamino acids, 1mM MgSO_4_; pH 7.0) were used to grow RP437-derived strains, AW405, and TW09231. H1 minimal salts medium (MMH1: 50 mM KPO_4_, 0.5 mM MgSO_4_, 7.6 mM (NH_4_)_2_SO_4_, 1.25 μM Fe_2_(SO_4_)_3_; pH 7.0) supplemented with 0.5% v/v glycerol and 0.01% w/v thiamine hydrochloride was used to grow MG1655-derived strains. MOPS glycerol media (8.37 g/L MOPS adjusted to pH 7.0 with KOH, 0.712 g/L tricine adjusted to pH 7.0 with KOH, 0.00278 g/L FeSO_4_ 7H_2_O, 0.0481 g/L K_2_SO_4_, 0.0000555 g/L CaCl_2_, 0.106 g/L MgCl_2·_6H_2_O, 2.91 g/L NaCl, 0.0230 g/L K_2_PO_4_, 2 mM NH_4_Cl, and 4 mL/L glycerol) were used to grow HE205 derived strains. To grow overnight cultures, small inoculants from the freezer stocks were added into test tubes containing 2 mL of media (with appropriate concentrations of inducers and antibiotics) in a shaking incubator at 30°C rotating at 250 RPM. To grow day cultures for cell tracking experiments, RP437-derived strain overnight cultures were diluted by 100-fold and re-grown in 5 mL of the appropriate media until exponential phase (OD_600_ = 0.20). For receptor quantification experiments, MG1655-derived strains were diluted 200-fold and re-grown in 10 mL of the appropriate media until exponential phase (OD_600_ = 0.30). For cell tracking experiments, chemotaxis buffer (1x M9 buffer, 0.4% v/v glycerol, 0.010 mM methionine, 0.1 mM EDTA, and 0.1% v/v PVP-40) was used to wash and dilute cells. For receptor quantification experiments, minimal chemotaxis buffer (10 mM KPO_4_ and 0.1mM EDTA; pH 7.0) was used to wash and dilute cells. For the TB measurements in batch cultures, RP437 *ΔcheZ pTet-cheZ* were grown in media supplemented with aTc that was aged for 15 hours.

### Growth rates measurements

Growth measurements were done using a microplate reader (BioTek Epoch 2 microplate spectrophotometer). Overnight cultures were diluted by 100-fold into fresh media supplemented with appropriate inducer concentrations. 200 μL of diluted cells were aliquoted into each well of polystyrene 96-well plates (Falcon 96-wells REF 353072), and the plates were loaded into the plate reader. The plates were incubated at 30°C while continuously shaking linearly at 567 cpm. The OD_600_ of each well was measured every 7 minutes for 36 hours. To extract the growth rates of each sample, the slopes of the linear region (at the mid-exponential growth phases) of the natural log of OD_600_ versus time (hours) curves were calculated.

### Formation of propagating waves and expansion speed measurements using swim plate and capillary tube assays

For the swim plate assay, 3 µL of cells in exponential phase were inoculated at the center of semi-solid agarose plates for RP437-derived strains or 10 *µL* of cells in exponential phase for MG1655-derived strains. For RP437-derived strains, the semi-solid agarose plates were made using M9-based media, as described above, supplemented with 0.14% agarose (American Bioanalytical Agarose GPG/LE) and appropriate concentrations of aTc. Similarly, for MG1655-derived strains, the semi-solid agarose plates were made using H1 minimal media, as described above, supplemented with 0.14% agarose. The plates were incubated at 30°C for 15 hours (RP437-derived strains) or at 33.5°C for 48 hours (MG1655-derived strains). The edges of the migrating colonies were picked using pipette tips and diluted into chemotaxis buffer (for RP437-derived strains) or minimal chemotaxis buffer (for MG1655-derived strains) for subsequent tracking and imaging analyses. To measure the expansion speeds of bacterial populations in swim plates, images were taken with a Canon DS126291 camera every 30 minutes. The diameters of the expansion rings as a function of time and expansion speeds were extracted from the images using a customized MATLAB code.

For the capillary tube assay, the protocol was adapted from previous work ^14^. Overnight cultures were inoculated 100-fold into 5 mL of media with appropriate inducers and grown at 30°C until OD_600_ = 0.2. Cells were concentrated to an OD_600_ = 6. A 12-inch capillary tube was filled with growth media (M9 glycerol with casamino acids, MgSO_4_, and PVP-40) and 6-hours aged aTc, and one side of the tube was plugged by stabbing the tube into a plate filled with solidified agar (1.5% agar in distilled water). The concentrated cell mixture was filled on the other side of the tube using a 1 mL syringe, and another agar plug was added into that side. Both sides of the tube were then plugged with clay. The tubes are incubated horizontally at 30°C for 9 hours. To isolate the wave, a 3-cm section of the tube that contained the wave was fractionated, and the mixture inside the 3-cm section was diluted into chemotaxis buffer for subsequent tracking analysis. To measure the expansion speeds of bacterial populations in capillary tubes, images were taken with a Canon DS126291 camera every 10 minutes. Positions of the populations as a function of time and expansion speeds were extracted from images using ImageJ.

### Microfluidic device preparation

To track single cell swimming behavior, we utilized a rectangular channel microfluidic device (0.5 mm wide, 30 mm long, and 60 *µ*m deep). The device was constructed from PDMS with standard lithography methods. To cast the device, a pre-prepared mold was coated with a 5-mm thick layer of degassed 10:1 PDMS-to-curing agent mixture (Sylgard 184, Dow Chemical). The mold was then baked at 70°C for 12 hours. Devices were cut out of the mold, and a hole was punched using a 20-gauge blunt-tip needle on each side of the channel. The device was rinsed with isopropanol, methanol, and water. A 22 mm x 50 mm glass coverslip was rinsed with acetone, isopropanol, methanol, and water. The coverslip and device were then treated inside a plasma cleaner for 1 minute and bonded together. The bonded device was heated at 80^c^C for 15 minutes and cooled to room temperature before cells were injected inside.

### Tracking single cells and tumble detection

Washed cells were diluted with chemotaxis buffer to an OD_600_ = 0.0005. The diluted mixture was injected inside the microfluidic device using a 1 mL syringe. To prevent evaporation, holes on both sides of the microfluidic device were taped. The device was placed on an inverted microscope (Nikon Eclipse Ti-U) equipped with a custom environmental chamber (50% humidity and 30°C). A custom MATLAB script was used to record 3-minute phase contrast videos at 4X magnification and 20 frames-per-second. Data from video recordings were stored as .bin files.

A customized MATLAB code was used to detect tumble and extract tumble bias of single cells, as previously described^58^. Only trajectories longer than 10 seconds were considered for analysis. Each tumble bias distribution was generated using 4 3-minute videos that contained ∼1000 total trajectories.

### Measuring the time scales of relaxation of TB distribution to the standing batch distribution

*E. coli* RP437 or HE205 were grown to mid-exponential phases and let to migrate in capillary assays for 9 hours, as mentioned above. The sections of the capillary tubes containing the migrating bacterial populations were fractioned, and the cells were added into 1 mL of growth media. For every 15- or 30-minutes during growth in the batch cultures, the TB distributions of the populations were measured. The mean of TB distributions normalized by the trajectory lengths were determined for each time points.

The equation to model for the relaxation of mean TB is:

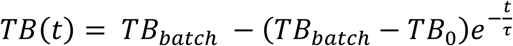

where *TB*(*t*) is the mean TB over time, *TB*_*batch*_ is the mean TB of the batch culture, *TB*_0_ is the mean TB during collective migration in the capillary tubes, and *τ* is the relaxation timescale. To estimate *τ*, we fitted the means of the mean TBs across replicates and the standard error of the means across replicates to the equation using maximum likelihood estimation. To estimate the error of *τ*, we sampled the posterior distribution using Markov Chain Monte Carlo simulation to get 100 simulated parameter values and calculate the standard deviation of those values.

### Protein expression measurements in swim plates

Non-motile MG1655-derived cells expressing labeled Tsr or non-motile RP437-derived cells expressing labeled CheY and CheZ used as biosensors for gene expression were grown as described above. Subsequently, cells were mixed with the appropriate media and 0.14% liquified agarose to a final OD_600_ = 0.01. Swim plates were left to solidify at room temperature for 2-3 hours, and afterward, 10 μL of OD_600_ = 0.01 motile WT MG1655 (labeled Tsr experiments) or motile WT RP437 (labeled CheY and CheZ experiments) cells were inoculated in the center of the plate. The motile population of cells shapes the gradient of the plate and ensures that the non-motile biosensor strain experiences the same local environment as motile cells. After a colony of migrating cells has formed, cells from the center and the edge of the colony are picked and diluted into minimal chemotaxis buffer for subsequent imaging.

### Single-cell protein copy number quantification using fluorescent microscopy

Washed cells were diluted 10-fold and plated on agarose pads (3% agarose in minimal chemotaxis buffer). The cell suspension was left to dry for 10 minutes, and cells were immediately imaged afterward. Imaging was performed using an inverted microscope (Nikon Eclipse Ti-E) equipped with an oil immersion 100x phase-contrast objective lens. To measure the fluorescence intensity of the strain expressing cheY-mRFP and cheZ-mYFP, fluorescent proteins were excited using a light-emitting diode illumination system (pE-4000, CoolLED). The RFP and YFP channels were imaged consecutively using a multi-band filter (Semrock), and the fluorescence emissions were led into a 2-camera image splitter (TwinCam, Cairn), leading to two identical sCMOS cameras (ORCA-Flash4.0 V2, Hamamatsu). The same setup was used to measure the fluorescence intensity of the strain expressing Tsr-mYFP. In a typical experiment, 20 fields of view, containing approximately 1000 cells in total, are imaged.

### Fluorescent image analysis for protein copy number quantification

For each field of view, one phase contrast, one YFP, and one RFP channel images were acquired. Using a custom MATLAB script, images from the two cameras were aligned using an affine transformation, and cells were segmented on the phase-contrast channel using a modified Otsu algorithm. Then, the image background was subtracted from each image, and the fluorescent intensity of every cell was extracted. Finally, the area of each cell was calculated by fitting an ellipsoid. The fluorescent intensity of each cell was divided by the cell area as a measure of protein concentration.

### Reproducibility and statistical analysis

All experiments reported in this work were repeated at least 3 times. All measurement values of TB, expansion speeds, growth rates, protein levels were reported as the average of the means of the distributions across biological replicates. Uncertainties in these average values were reported as the standard deviation of the means of the distributions.

To construct the TB distributions, TB values were sorted into bins, and the probability density were normalized by dividing the trajectory time within each bin by the total trajectory time. To represent the variations in TB distributions, the standard error of the mean TBs within each bin was calculated. Similarly, with the Tsr intensity distribution, single-cell Tsr intensities were sorted into bins, and the probability density in each bin was calculated. The variations in Tsr intensity distributions are represented as the standard error of the mean intensities within each bin.

To construct the fold-enrichment curve, the probability density values of each TB or Tsr intensity bins observed in the distributions during collective migration were divided by the values observed in batch cultures. Variations in fold-enrichments are represented as the standard error of the mean fold-enrichment values within each bin.

To determine statistical significance between TB and protein level distributions, two-sided t-test was performed on the mean values. Similarly, two-sided t-test was performed to determine significance of growth rates and expansion speeds between experiments (throughout, *** = P<0.0005, ** = P<0.005, * = P<0.05, and n.s. = not significant).

## Analysis code and data availability

Analysis code will be uploaded to the Emonet lab GitHub (github.com/emonetlab) upon publication. Data will be made available on datadryad.org upon publication.

## Acknowledgments

We thank Jing Yan, Amir Pahlavan, Jeremy Moore, Jyot Antani, Karen Fahrner, Kiri Choi, and Gustavo Santana for useful discussions. We thank Simone Boskamp for assistance with cloning and Cherin Mohamed for assistance with experiments. We thank Jonas Cremer and Terrence Hwa for providing *E. coli* HE205 wild-type strain. We thank Victor Sourjik and Howard Berg for providing plasmids and strains that were important for this work.

## Author Contributions

L.V., F.A., H.M., T.S.S, B.I.K., and T.E. designed the project and experiments. L.V., F.A., K.E., I.B., and R.B. performed the experiments. L.V. and F.A. performed the analyses. L.V., F.A., H.M., T.S.S, B.I.K., and T.E. wrote the manuscript.

## Supporting Information for

**Figure S1:**
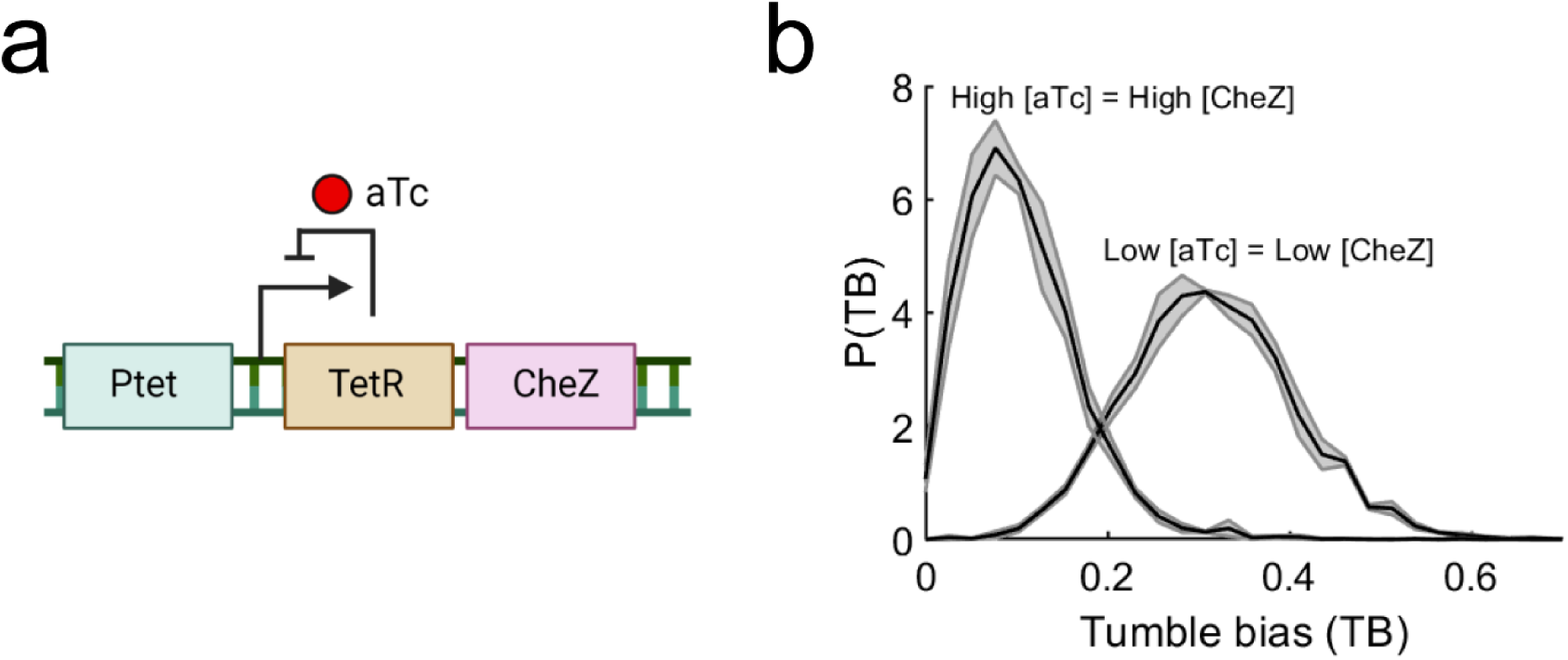
A genetic construct that tunes TB distribution of a population. **(a)** To tune tumble bias distributions, we utilized an *E. coli* RP437 *ΔcheZ* strain that carries an aTc-inducible cassette of *cheZ* that is integrated in the chromosome at the *attB* site (RP437 *ΔcheZ pTet-cheZ*). **(b)** *E. coli* populations with different initial TB distributions were generated by inducing RP437 *ΔcheZ pTet-cheZ* strain with different aTc concentrations. High aTc concentration (16 ng/mL) formed low mean TB populations (mean TB = 0.10 ± 0.01; *N*_*rep*_ = 3, *N*_*cells*_ = 21,891), and low aTc concentration (2 ng/mL) formed high mean TB populations (mean TB = 0.31 ± 0.01; *N*_*rep*_ = 3, *N*_*cells*_ = 26,254).

**Figure S2:**
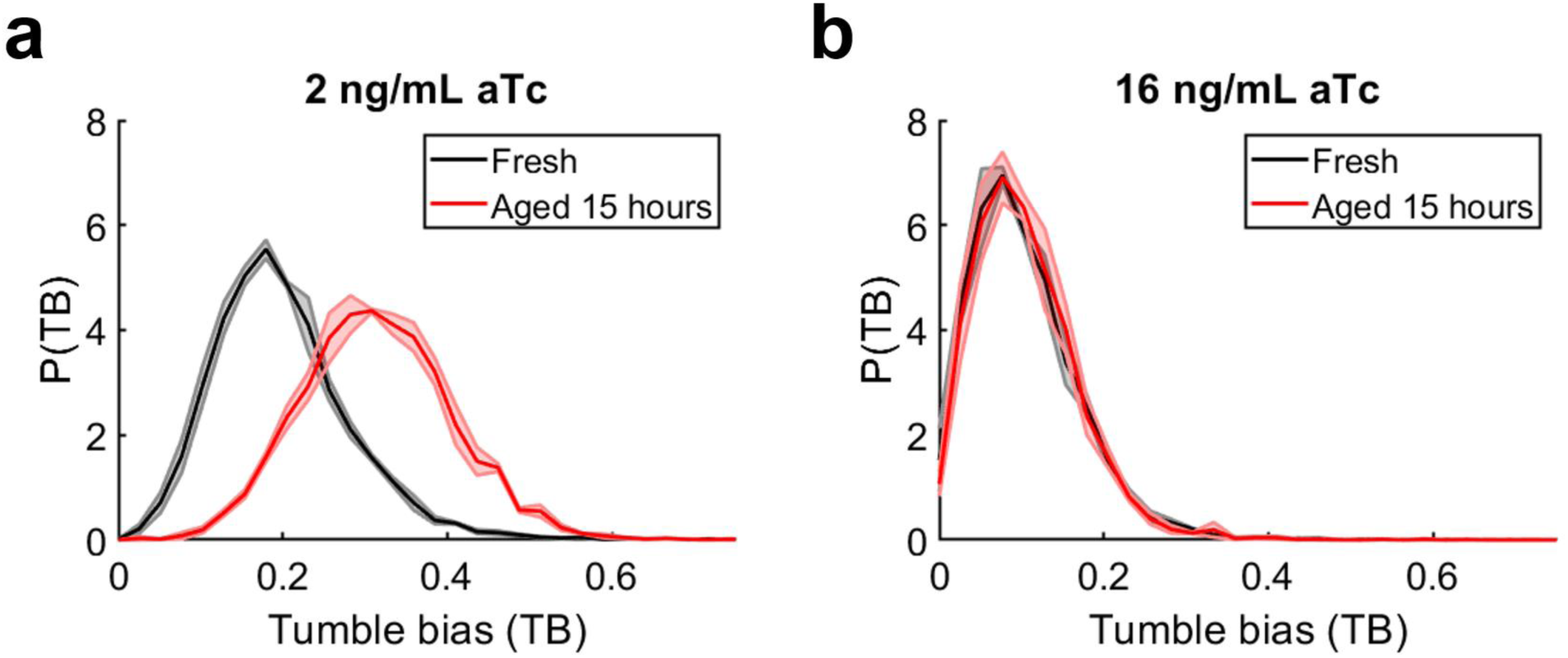
aTc degrades over time. Tumble bias distribution of RP437 *ΔcheZ pTet-cheZ* grown in batch culture supplemented media with 2 ng/mL (mean TB = 0.2 ± 0.01; *N*_*rep*_ = 3, *N*_*cells*_ = 42,301), **(a)** and 16 ng/mL (mean TB = 0.10 ± 0.01; *N*_*rep*_ = 3, *N*_*cells*_ = 21,093), **(b)** of fresh aTc or aTc that was aged for 15 hours (same as in **Fig. 1b**). 15 hours was how long we waited until we isolated the edge of the migrating colony. Except for 16 ng/mL aTc, the distributions of tumble biases increased in mean values if the strain was grown in aged aTc, suggesting that aTc degrades in the growth media over time.

**Figure S3:**
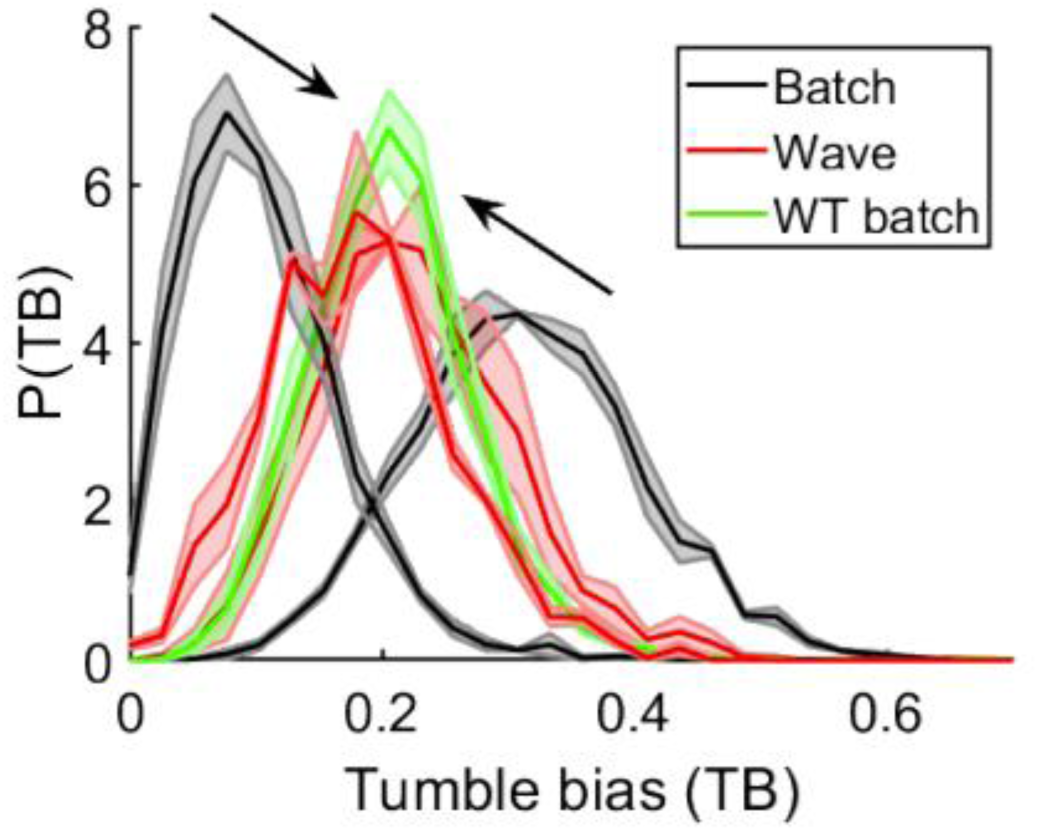
Migration in agarose swim plate tunes the population’s TB distribution toward the TB distribution of wild-type population. The black curves are the TB distributions of low- and high-TB populations grown in batch culture, and the red curves are the TB distributions of those population after migration on agarose swim plates, as shown in **Fig. 2c** and **e**. The green curve is the TB distribution of wild-type *E. coli* RP437 grown in batch culture (mean TB = 0.22 ± 0.03; *N*_*rep*_ = 3, *N*_*cells*_ = 9,417).

**Figure S4:**
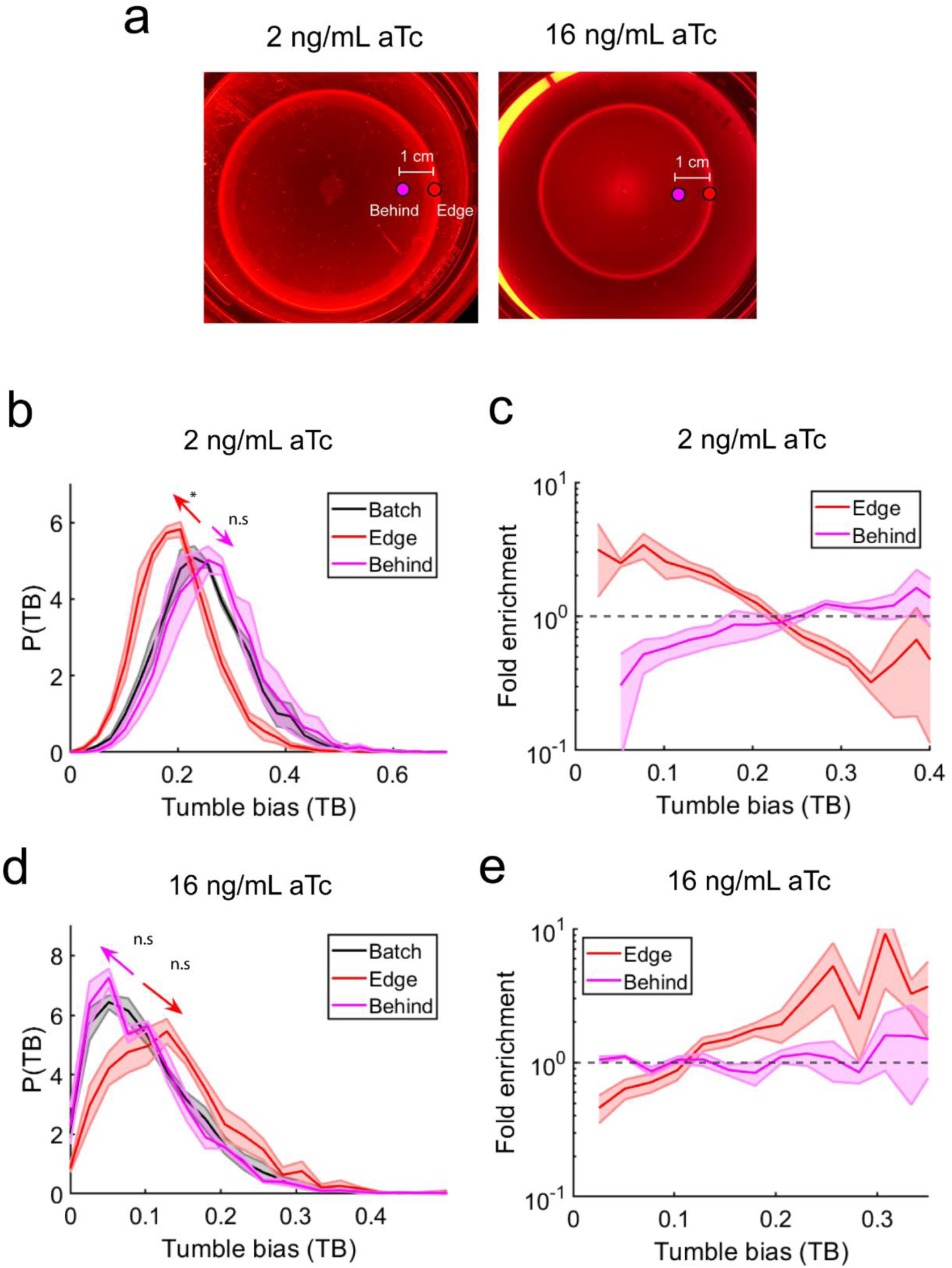
TB distributions of cell populations that fall behind during collective migration. (**a**) Experimental design: Low-TB and high-TB *E. coli* HE205 Δ*cheZ* (pTet-CheZ) populations (“batch”) was inoculated on swim plates supplemented with 100 μM aspartate and appropriate inducer concentrations. After 16 hours of growth and migration on swim plates, the TB distributions of populations picked at the edge (red) and middle (located ∼1 cm behind the edge, magenta) were measured. (**b**) Populations with higher initial mean TB (0.25 ± 0.01; *N*_*rep*_ = 3, *N*_*cells*_ = 5,164) shift toward a lower mean TB after migration on the plates (0.20 ± 0.01; *N*_*rep*_ = 3, *N*_*cells*_ = 6,307), similar to our observation in E. coli RP437 (see **Fig. 2e**). The TB distributions of populations at the “middle” (mean TB = 0.26 ± 0.02; *N*_*rep*_ = 3, *N*_*cells*_ = 5,602) were similar to the TB distributions observed in batch cultures, and the TB distributions of populations at the “center” have the highest mean TB. (**c**) Cells with low TBs were enriched, while cells with high TBs were filtered at the edge of the migrating colonies. Cells with low TBs were depleted behind the wave (*N*_*rep*_ = 3). (**d**) Same as (**b**) but with populations with lower initial mean TB. The TB distributions of populations with lower mean TB (0.10 ± 0.01; *N*_*rep*_ = 3, *N*_*cells*_ = 2,090) shifted to a higher mean TB value (0.13 ± 0.01; *N*_*rep*_ = 3, *N*_*cells*_ = 3,151) during migration on the swim plates. The population that fell behind exhibited similar TB distributions to the batch cultures (0.10 ± 0.01; *N*_*rep*_ = 3, *N*_*cells*_ = 2,625). Shaded area is the standard errors of the probability density values at each TBs ( *N*_*rep*_ = 3, *N*_*cells*_ > 1000 for each TB distributions). (**e**) Cells with higher TBs were enriched, while cells with very low TBs were filtered out at the edge of the migrating colonies. No enrichments were observed behind the migrating edge (*N*_*rep*_ = 3). Two-sided T-tests were performed to determine significance of the difference between pairs of means throughout (*** = P<0.0005, ** = P<0.005, * = P<0.05, and n.s. = not significant).

**Figure S5:**
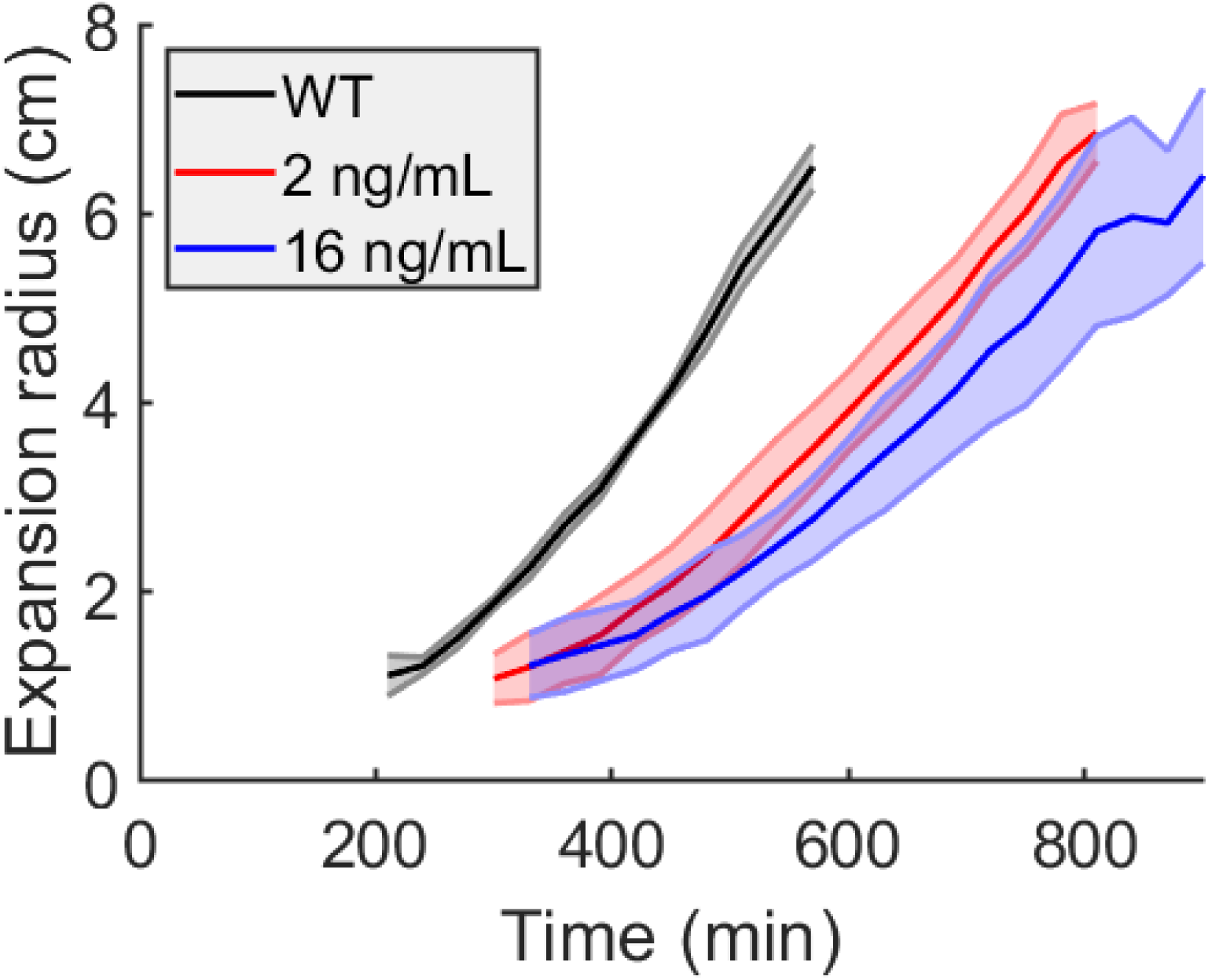
Expansion dynamics of migrating *E. coli* RP437 populations on swim plates. Expansion radius of wild-type *E. coli* RP437 and the CheZ-inducible population (induced with 2 ng/mL and 16 ng/mL aTc) over time. Shaded area is the standard deviation of the mean expansion radius and speed. The curves start at when the expansion ring became visible on the swim plate assays (*N*_*rep*_ = 6 for the 2 ng/mL and 16 ng/mL curves; *N*_*rep*_ = 12 for the WT curve).

**Figure S6:**
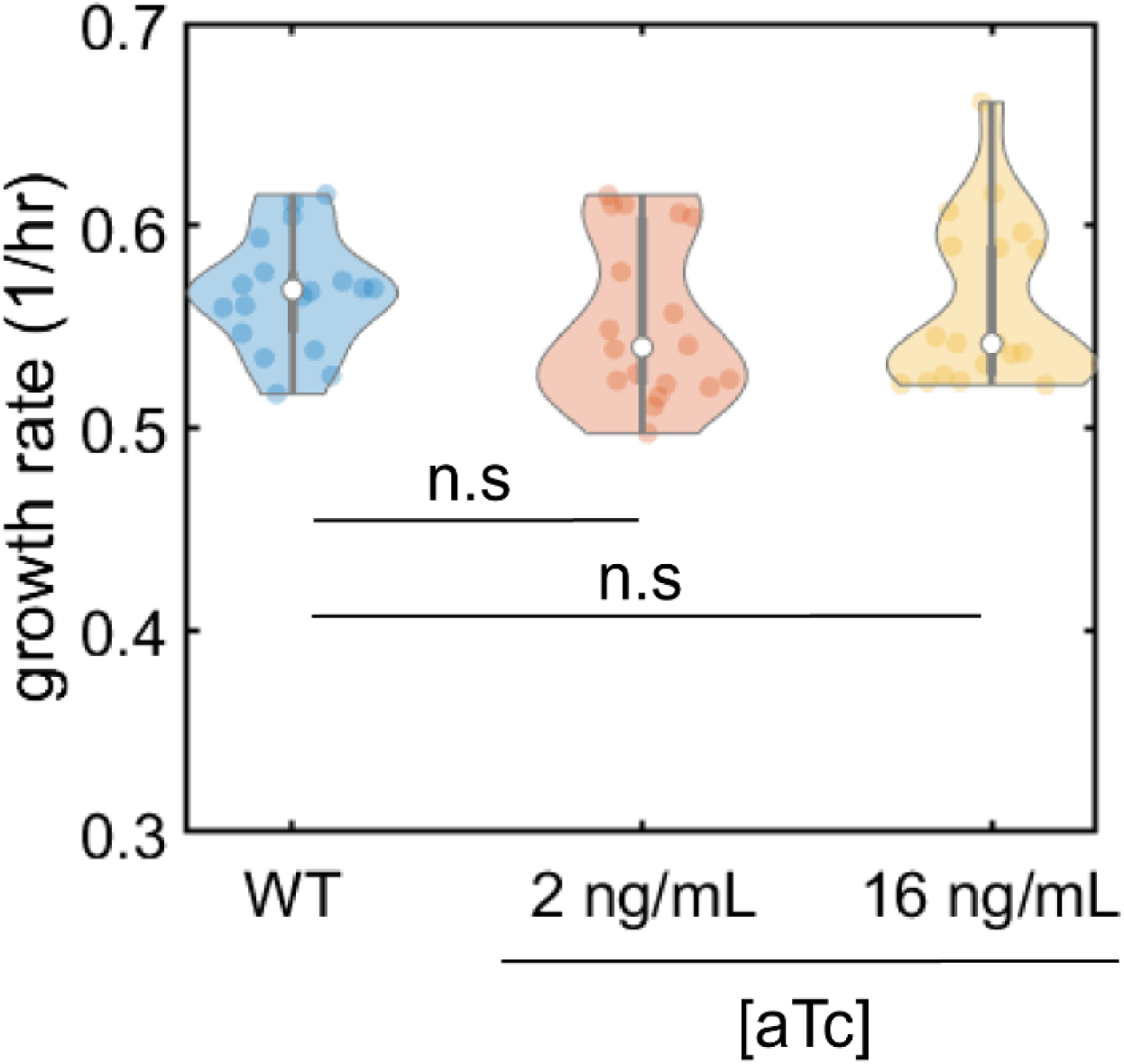
The growth rates of RP437 wild-type and *ΔcheZ* (pTet-CheZ) populations are identical. Growth rates of RP437 wild-type (mean 0.5663 ± 0.03 1/hr, throughout, uncertainty is the standard deviation of the means across replicates, *N*_*rep*_ = 3), *ΔcheZ* (pTet-CheZ) induced with 2 ng/mL aTc (mean 0.5526 ± 0.04 1/hr, *N*_*rep*_ = 3), and 16 ng/mL aTc (mean 0.5608 ± 0.04 1/hr, *N*_*rep*_ = 3) populations. The differences in growth rates cannot explain the differences in the expansion speeds of WT and *ΔcheZ* (pTet-CheZ) populations in agarose swim plates and capillary tube assays. Throughout, two-sided T-tests were performed to determine statistical significance of the difference between pairs of means (*** = P<0.0005, ** = P<0.005, * = P<0.05, and n.s. = not significant).

**Figure S7:**
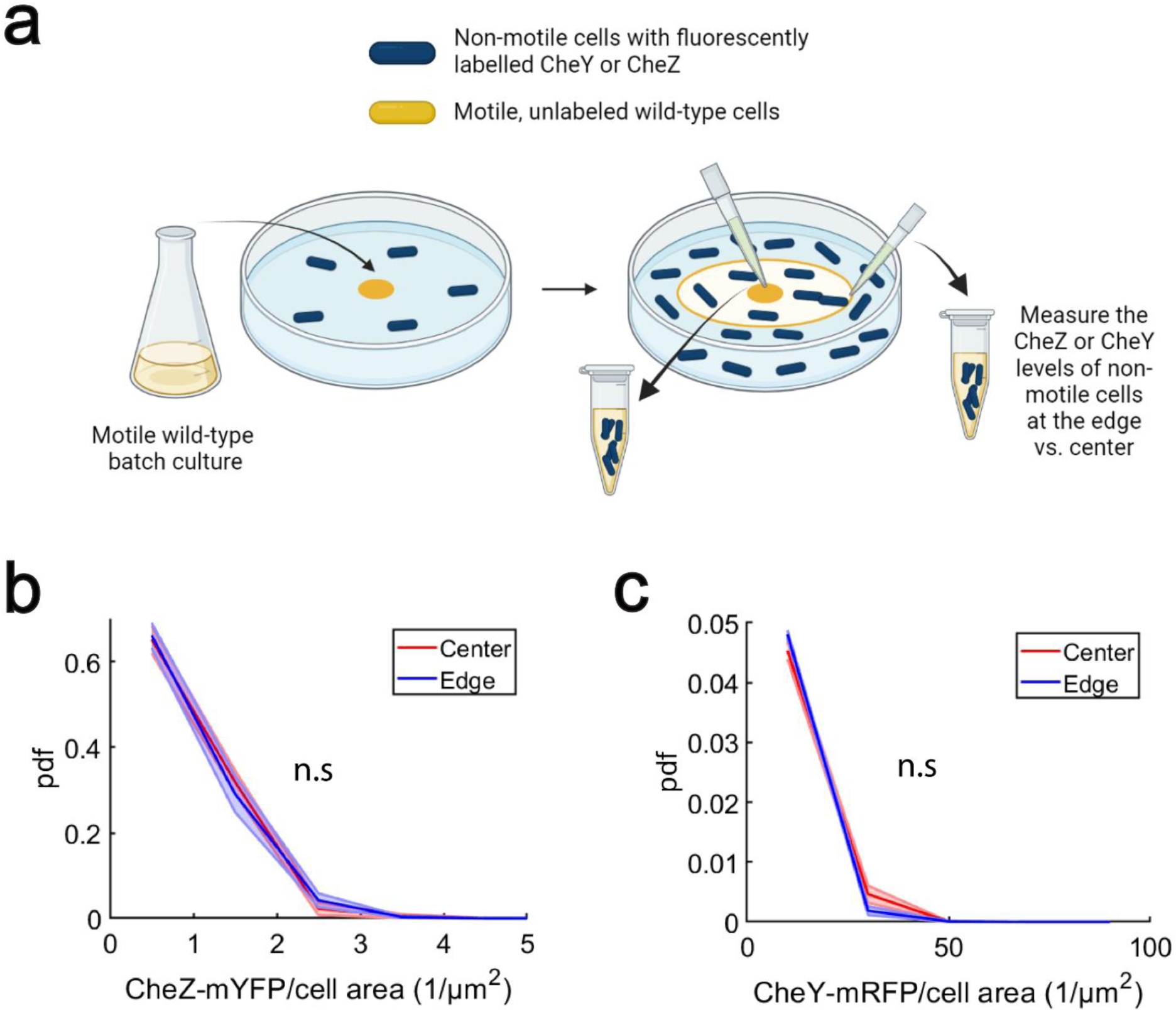
The shift in TB distribution is not due changes in the CheY and CheZ expressions. **(a)** Motile RP437 wild-type bacteria migrated in agarose swim plates that were filled homogenously with non-motile *E. coli* RP437 cells with labelled CheY or CheZ (CheY-mRFP or CheZ-mYFP respectively). Fluorescent signal of CheY or CheZ of the biosensor were measured at the center versus the edge of the migrating colonies. **(b)** Distribution of CheZ-mYFP over a single cell’s area at the center (mean CheZ intensity = 0.96 ± 0.02 *µm*^−2^; *N*_*rep*_ = 3, *N*_*cells*_ = 739) and the edge (mean CheZ intensity = 6.3 ± 4.6 *µm*^−2^; *N*_*rep*_ = 3, *N*_*cells*_ = 1,076) of the colony. **(c)** Distribution of CheY-mRFP over a single cell’s area at the center (mean CheY intensity = 0.95 ± 0.19 *µm*^−2^; *N*_*rep*_ = 3, *N*_*cells*_ = 739) and the edge (mean CheY intensity = 11.3 ± 1.4 *µm*^−2^; *N*_*rep*_ = 3, *N*_*cells*_ = 1,076) of the colony. The distributions at the center and edge of the migrating colonies are similar, suggesting that the shift in TB distributions observed after migration on agarose swim plates is not due to changes in gene expression of CheY and CheZ. Shaded area is the standard error of the mean probabilities within each bin of CheZ or CheY intensity values. Two-sided T-tests were performed to determine significance of the difference between pairs of means (*** = P<0.0005, ** = P<0.005, * = P<0.05, and n.s. = not significant).

**Figure S8:**
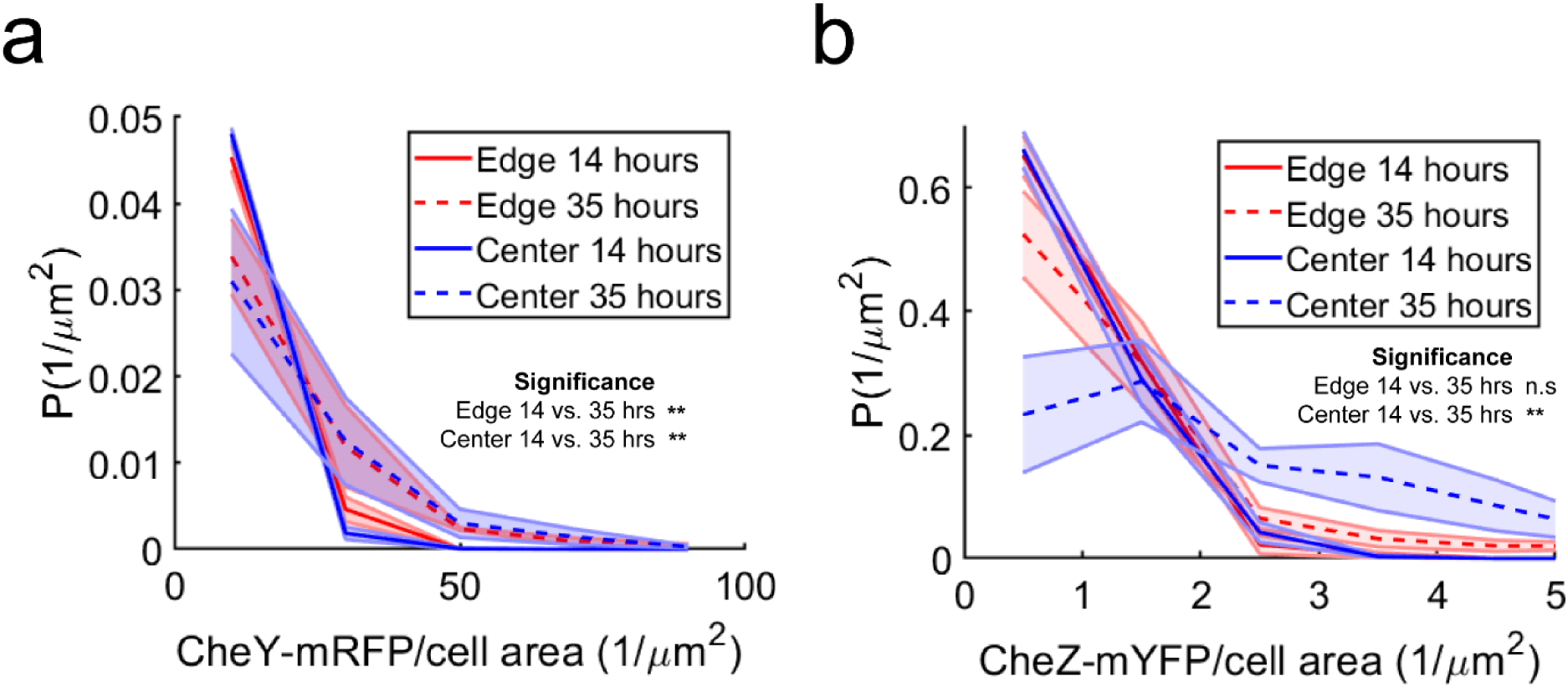
Cells experience catabolite repression after prolonged migration. Using the same experimental setup (shown in **Fig. S7**), the CheY and CheZ expressions were measured at the center versus the edge of the migrating colonies at 14 hours and 35 hours. After 35 hours of migration, populations at the edge and center exhibit higher CheY (**a**) and CheZ (**b**) expressions. The CheY expression of populations isolated at the center (mean CheY intensity = 22.3 ± 10.2 *µm*^−2^; *N*_*rep*_ = 3, *N*_*cells*_ = 1,090) and edge (18.5 ± 2.72 *µm*^−2^; *N*_*rep*_ = 3, *N*_*cells*_ = 1,993) are not significantly different. The CheZ expression of populations picked at the center (mean CheZ intensity = 2.67 ± 0.65 *µm*^−2^; *N*_*rep*_ = 3, *N*_*cells*_ = 1,090) is significantly higher than the CheZ expression of populations picked at the edge (1.43 ± 0.08 *µm*^−2^; *N*_*rep*_ = 3, *N*_*cells*_ = 1,993). Shaded area is the standard errors of the probability density values of CheY and CheZ expressions (*N*_*rep*_ = 3, *N*_*cells*_ > 600 for each distribution). Two-sided T-tests were performed to determine significance of the difference between pairs of means of CheY and CheZ expressions.

**Figure S9:**
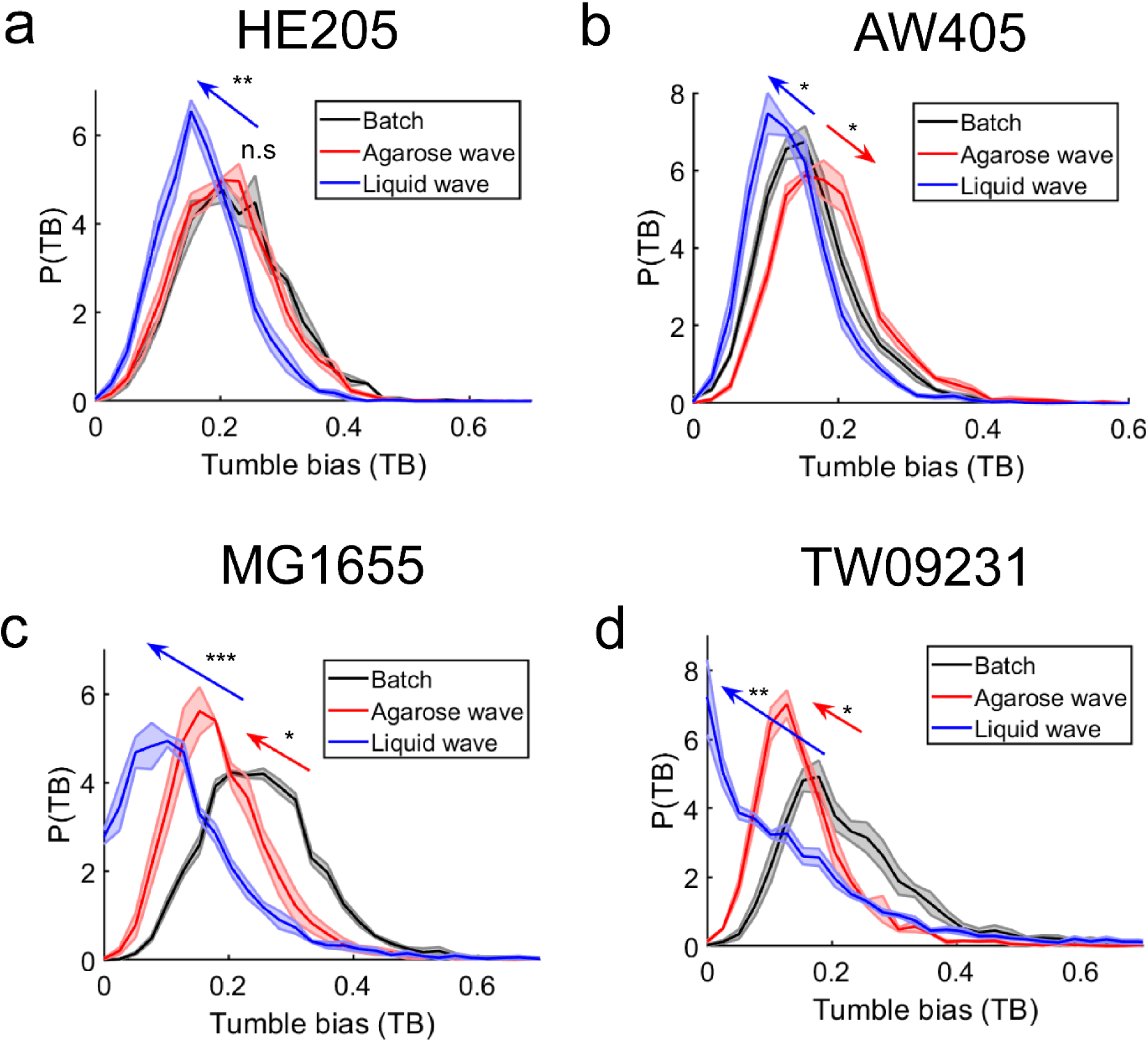
Environment-dependent tuning of TB distributions across *E. coli* strains. TB distributions of *E. coli* HE205, AW405, MG1655, and TW09231 (*Escherichia sp.* TW09231) were measured during collective migration in agarose swim plates, capillary tube assays, and during growth in batch cultures. (**a**) The mean TB values of HE205 during migration in agarose swim plates (mean TB = 0.21 ± 0.01; *N*_*rep*_ = 3, *N*_*cells*_ = 3,134) were the same as the mean TB values observed in batch cultures (0.22 ± 0.005; *N*_*rep*_ = 3, *N*_*cells*_ = 5,164). On the other hand, the mean TB values were lowered during migration in capillary tube assays (0.17 ± 0.01; *N*_*rep*_ = 3, *N*_*cells*_ = 2,164). (**b**) The mean TB values of AW405 during migration in agarose swim plates (0.19 ± 0.004; *N*_*rep*_ = 3, *N*_*cells*_ = 4,634) were higher than the mean TB values observed in batch cultures (0.16 ± 0.003; *N*_*rep*_ = 3, *N*_*cells*_ = 3,038). The mean TB values were lowered during migration in capillary tube assays (0.14 ± 0.01; *N*_*rep*_ = 3, *N*_*cells*_ = 5,300). (**c**) The mean TB values of MG1655 during migration in agarose swim plates (0.19 ± 0.01 *N*_*rep*_ = 3, *N*_*cells*_ = 5,164) were significantly higher than the mean TB values observed in batch cultures (0.26 ± 0.004; *N*_*rep*_ = 3, *N*_*cells*_ = 5,916). The mean TB values were lowered during migration in capillary tube assays (0.16 ± 0.01; *N*_*rep*_ = 3, *N*_*cells*_ = 3,164). (**d**) The mean TB values of TW09231 during migration in agarose swim plates (0.16 ± 0.01; *N*_*rep*_ = 3, *N*_*cells*_ = 4,781) were significantly higher than the mean TB values observed in batch cultures (0.23 ± 0.02; *N*_*rep*_ = 4, *N*_*cells*_ = 3,497). The mean TB values were lowered during migration in capillary tube assays (0.14 ± 0.01; *N*_*rep*_ = 4, *N*_*cells*_ = 5,140). Two-sided T-tests were performed to determine significance of the difference between pairs of means (*** = P<0.0005, ** = P<0.005, * = P<0.05, and n.s. = not significant).

**Figure S10:**
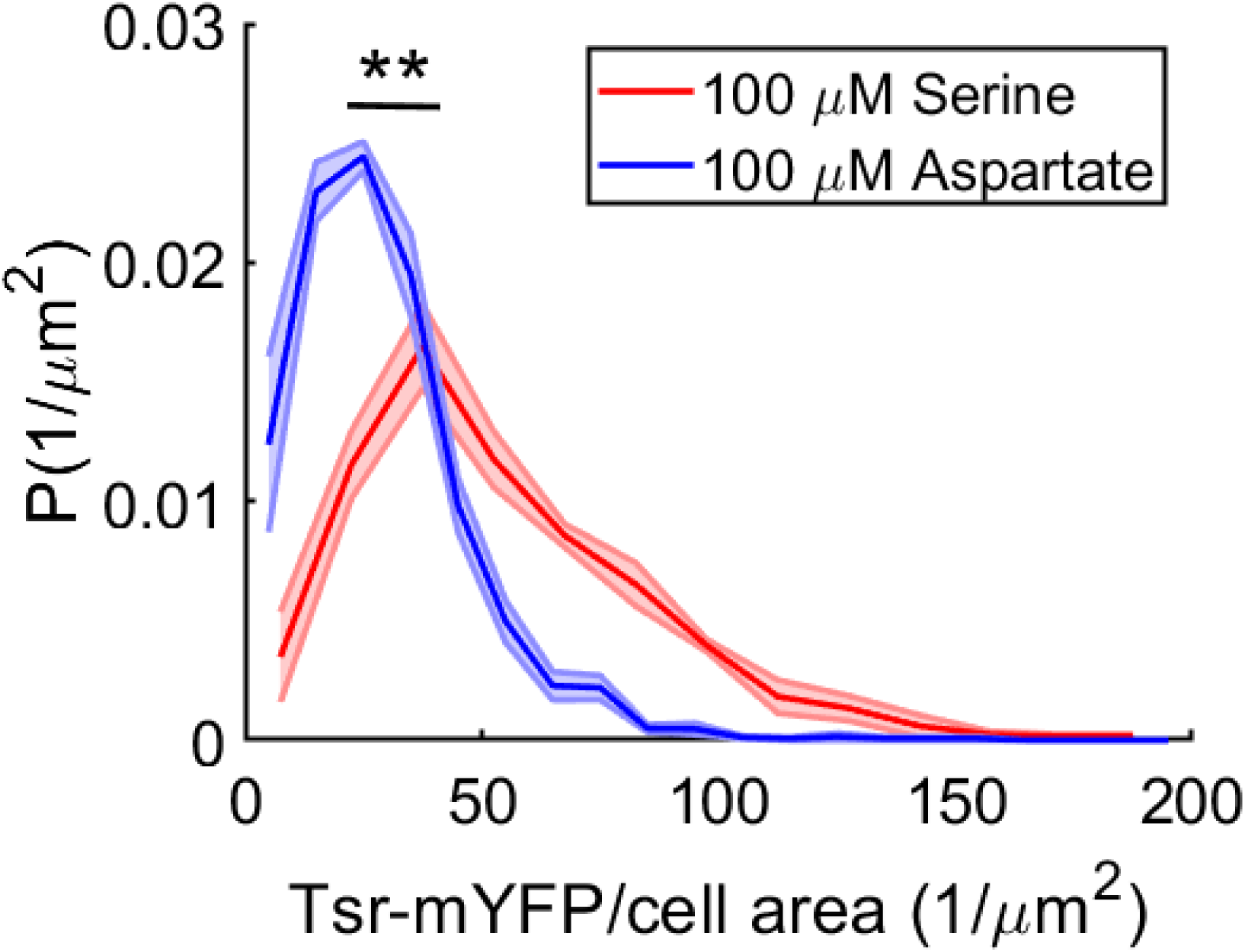
Direct comparison of the Tsr distributions during collective migration in serine and aspartate. The distributions of Tsr levels in populations of *E. coli* MG1655 wild-type with YFP-labelled Tsr when chasing 100 μM aspartate (mean Tsr intensity = 27.2 ± 3.8 *µm*^−2^; *N*_*rep*_ = 3, *N*_*cells*_ = 1,617) or 100 μM serine (53.6 ± 6.5 *µm*^−2^; *N*_*rep*_ = 3, *N*_*cells*_ = 675) on swim plates. Shaded area is the standard errors of the probability density values of Tsr expressions ( *N*_*rep*_ = 3, the distributions are the same as in **Fig. 6b** and **d**). Two-sided T-tests were performed to determine significance of the difference between pairs of means of Tsr intensities (*** = P<0.0005, ** = P<0.005, * = P<0.05, and n.s. = not significant).

**Figure S11:**
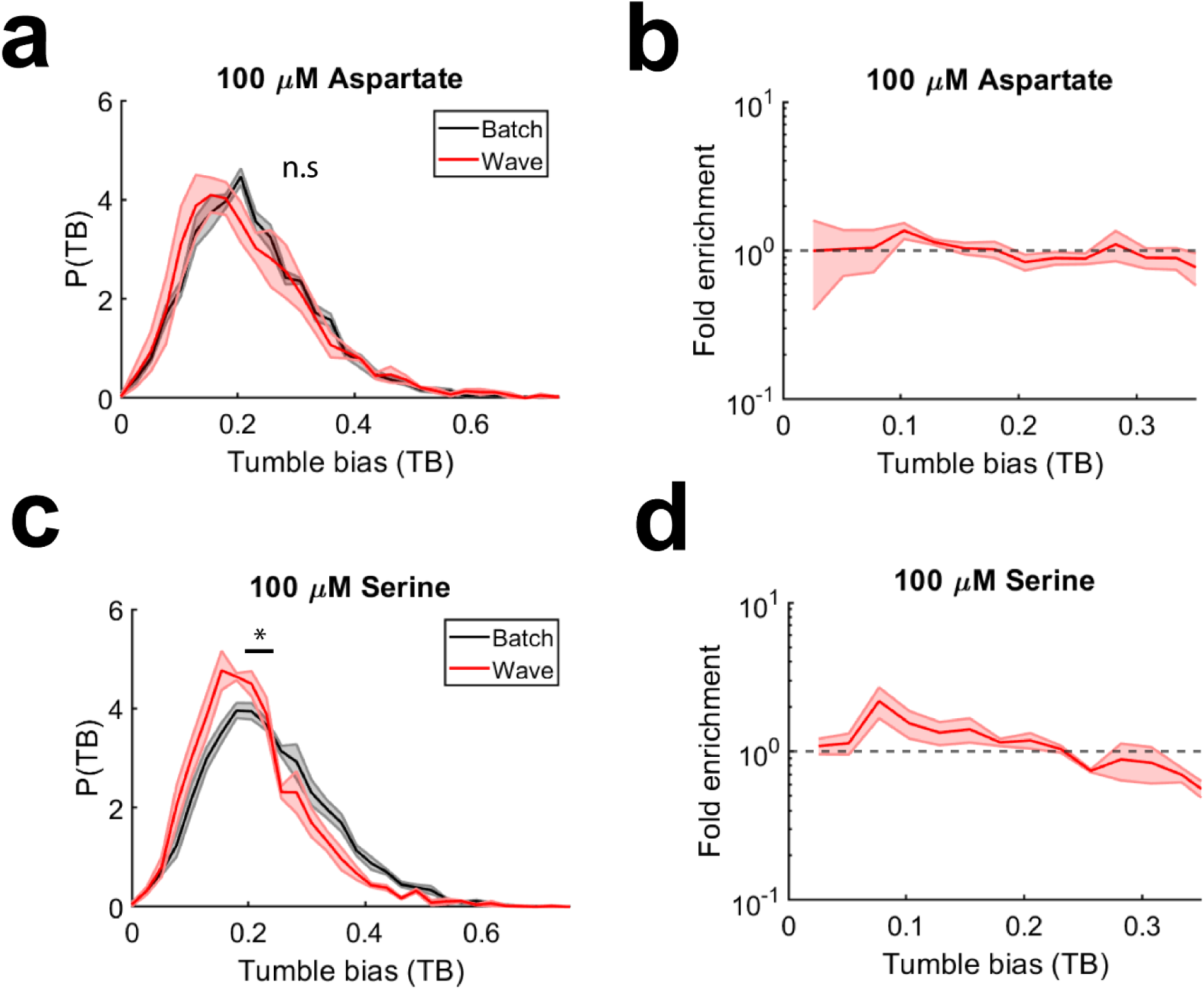
Tuning of TB distributions when the migrating populations are chasing aspartate and serine. Motile *E. coli* MG1655 cells expressing Tsr-YFP was grown in agarose swim plate supplemented with 100 μM aspartate **(a,b)** or 100 μM serine **(c,d)**. The TB distributions of populations isolated from the edge of the migrating colonies were measured and compared to the TB distributions observed in batch cultures, which were also supplemented with aspartate or serine. There was no significant differences between the mean TB of populations grown in batch cultures (mean TB = 0.23 ± 0.004; *N*_*rep*_ = 3, *N*_*cells*_ = 9,321) and the mean TB of populations migrated in swim plates supplemented with aspartate (0.22 ± 0.02; *N*_*rep*_ = 3 *N*_*cells*_ = 4,361). On the other hand, the mean TB of populations migrated in swim plates supplemented with serine (0.21 ± 0.01; *N*_*rep*_ = 3, *N*_*cells*_ = 3,686) exhibited a slightly, but significantly lower mean TB values compared to the mean TB value observed in batch cultures (0.25 ± 0.01; *N*_*rep*_ = 3, *N*_*cells*_ = 7,888). **(b,d)** Fold-enrichment of TB during collective migration on swim plates. Migrating on swim plates supplemented with serine mildly, but statistically significantly enriches for lower TBs. Shaded area is the standard errors of the probability density values at each TB. Two-sided T-tests were performed to determine significance of the difference between pairs of means (*** = P<0.0005, ** = P<0.005, * = P<0.05, and n.s. = not significant).

**Figure S12:**
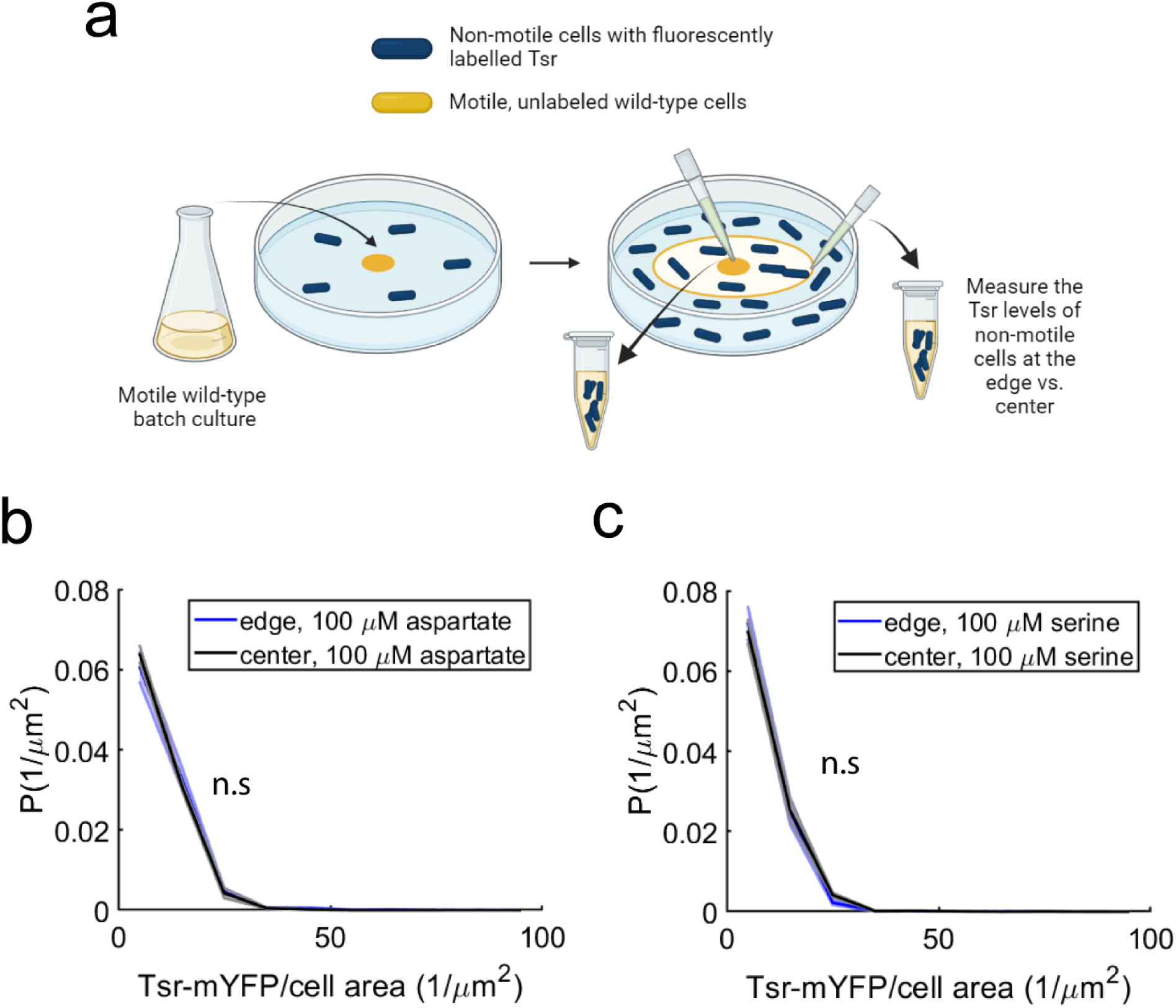
The shift in Tsr levels during collective migration is not due *tsr* regulation. **(a)** Motile MG1655 wild-type bacteria migrated in agarose swim plates (supplemented with 100 M Aspartate) that were filled homogenously with non-motile *E. coli* MG1655 cells expressing Tsr-YFP from the native locus, effectively acting as a biosensor for the regulation of gene expression by the passage of the migrating wave. Fluorescent signals, a proxy for *tsr* expression of the biosensor strain, were measured at the center versus the edge of the migrating colonies. **(b)** Distribution of Tsr levels at the center (mean Tsr intensity = 8.8 ± 0.5 *µm*^−2^; *N*_*rep*_ = 3, *N*_*cells*_ = 2,285) and the edge (mean Tsr intensity = 9.4 ± 1.0 *µm*^−2^; *N*_*rep*_ = 3, *N*_*cells*_ = 2,390) of the colony on plates supplemented with aspartate were identical indicating that there was no regulation of *tsr* expression upon passage of the migrating wave. (c) Center (mean Tsr intensity = 7.8 ± 0.6 *µm*^−2^; *N*_*rep*_ = 3, *N*_*cells*_ = 1911) and edge (mean Tsr intensity = 7.9 ± 1.0 *µm*^−2^; *N*_*rep*_ = 3, *N*_*cells*_ = 2696) same as b, but on plates supplemented with Serine. Shaded area is the standard error of the mean probabilities within each bin of Tsr intensity values. Two-sided T-tests were performed to determine significance of the difference between pairs of means (*** = P<0.0005, ** = P<0.005, * = P<0.05, and n.s. = not significant).

**Figure S13:**
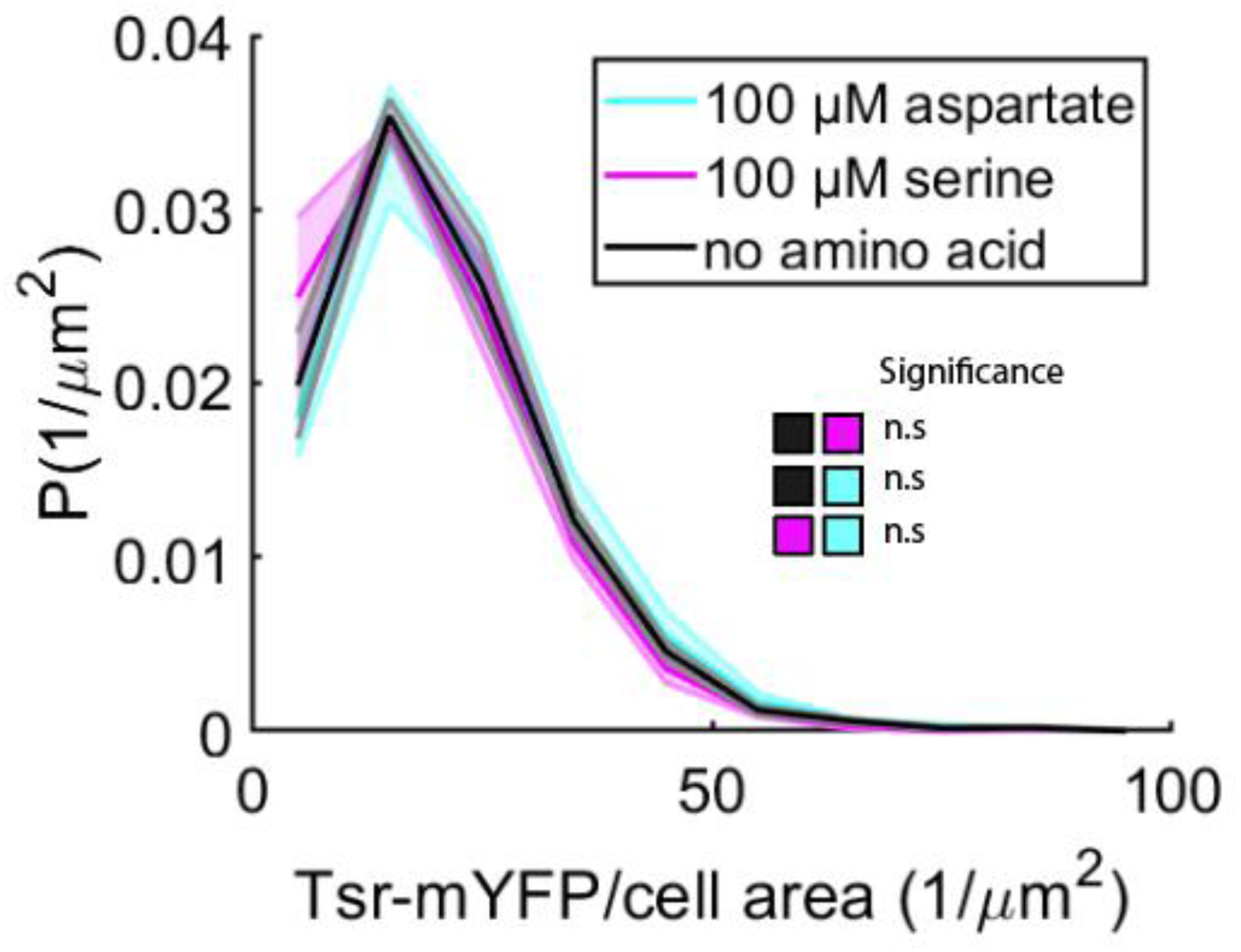
Expression of *tsr* is not regulated by serine or aspartate consumption during growth. Fluorescence of the biosensor *E. coli* MG1655 cells expressing Tsr-YFP from the native locus grown in batch culture with minimal media H1 supplemented either with no amino acids or with 100 μM serine or 100 μM aspartate were measured. Distribution of Tsr levels of cells when grown in serine (mean Tsr intensity = 18.7 ± 2.1 *µm*^−2^; *N*_*rep*_ = 3, *N*_*cells*_ = 1,695) in aspartate (mean Tsr intensity = 21.2 ± 2.5 *µm*^−2^; *N*_*rep*_ = 3, *N*_*cells*_ = 3,442), and with no amino acid added (mean Tsr intensity = 20.3 ± 1.7 *µm*^−2^; *N*_*rep*_ = 3, *N*_*cells*_ = 2,908). The distributions of Tsr levels are identical across conditions, suggesting that *tsr* expression is not regulated by aspartate or serine consumption. Shaded area is the standard error of the mean probabilities within each bin of Tsr intensity values. Two-sided T-tests were performed to determine significance of the difference between pairs of means (*** = P<0.0005, ** = P<0.005, * = P<0.05, and n.s. = not significant).

**Figure S14:**
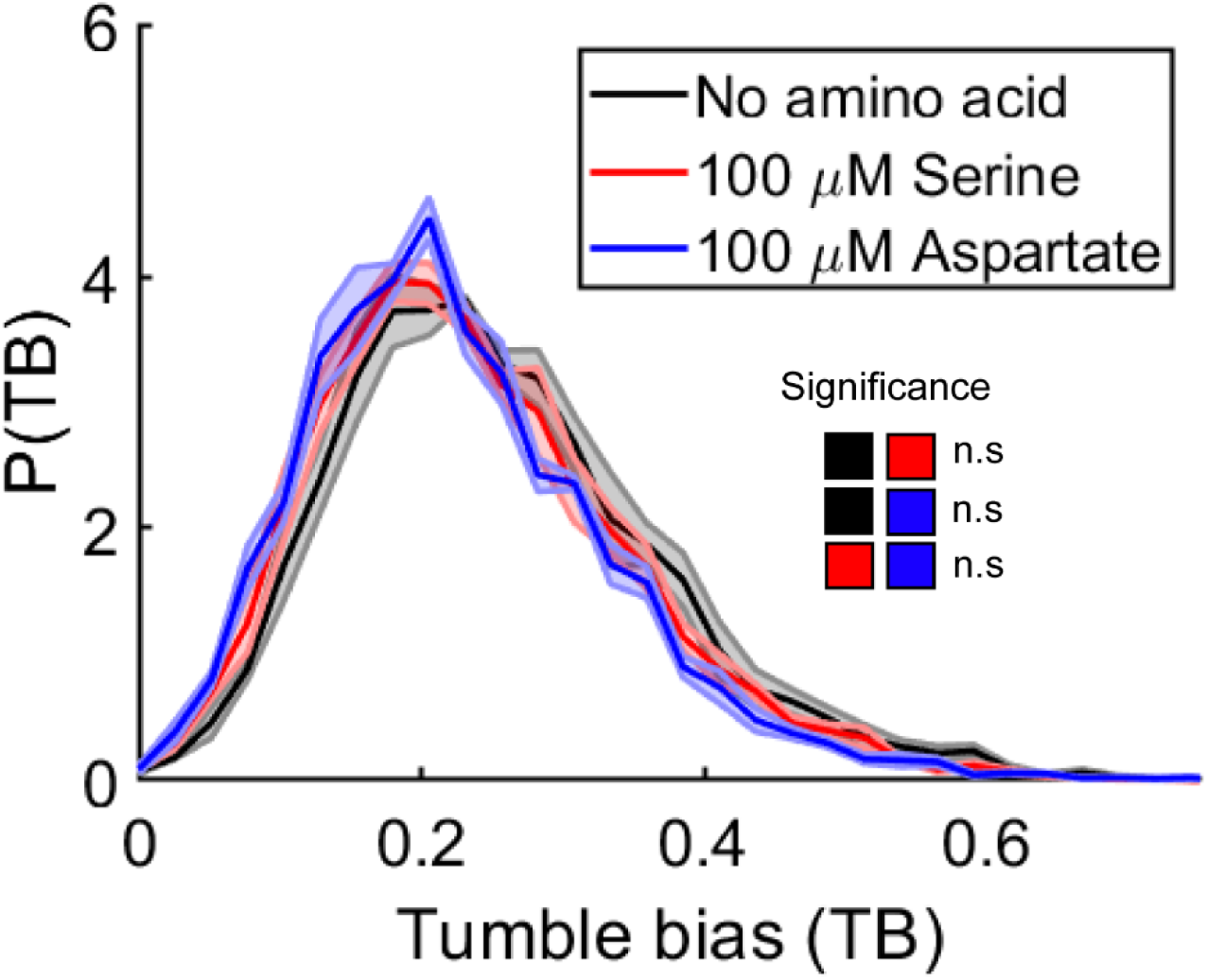
The presence of an attractant does not affect TB distributions. The TB distributions of motile *E. coli* MG1655 cells expressing Tsr-YFP were measured after growth in batch cultures supplemented with no amino acids (mean TB= 0.25 ± 0.01; *N*_*rep*_ = 3, *N*_*cells*_ = 6,111), 100 μM aspartate (0.23 ± 0.01; *N*_*rep*_ = 5, *N*_*cells*_ = 9,321), or 100 μM serine (0.25 ± 0.01; *N*_*rep*_ = 5 *N*_*cells*_ = 7,888). Addition of aspartate or serine did not significantly change the TB distributions of the populations. Shaded area is the standard errors of the probability density values at each TB bins. Two-sided T-tests were performed to determine significance of the difference between pairs of means (*** = P<0.0005, ** = P<0.005, * = P<0.05, and n.s. = not significant).

## References

1 Chan, B. K. et al. Phage selection restores antibiotic sensitivity in MDR Pseudomonas aeruginosa. Sci Rep 6, 26717 (2016). 10.1038/srep26717

2 Poole, K. Outer membranes and efflux: the path to multidrug resistance in Gram-negative bacteria. Curr Pharm Biotechnol 3, 77–98 (2002). 10.2174/1389201023378454

3 Schmitt, M. W., Loeb, L. A. & Salk, J. J. The influence of subclonal resistance mutations on targeted cancer therapy. Nat Rev Clin Oncol 13, 335–347 (2016). 10.1038/nrclinonc.2015.175

4 Liu, W., Cremer, J., Li, D., Hwa, T. & Liu, C. An evolutionarily stable strategy to colonize spatially extended habitats. Nature 575, 664–668 (2019). 10.1038/s41586-019-1734-x

5 Fraebel, D. T. et al. Environment determines evolutionary trajectory in a constrained phenotypic space. Elife 6 (2017). 10.7554/eLife.24669

6 Ni, B. et al. Evolutionary Remodeling of Bacterial Motility Checkpoint Control. Cell Rep 18, 866–877 (2017). 10.1016/j.celrep.2016.12.088

7 Norman, T. M., Lord, N. D., Paulsson, J. & Losick, R. Stochastic Switching of Cell Fate in Microbes. Annu Rev Microbiol 69, 381–403 (2015). 10.1146/annurev-micro-091213-112852

8 Kussell, E. & Leibler, S. Phenotypic diversity, population growth, and information in fluctuating environments. Science 309, 2075–2078 (2005). 10.1126/science.1114383

9 Balaban, N. Q., Merrin, J., Chait, R., Kowalik, L. & Leibler, S. Bacterial persistence as a phenotypic switch. Science 305, 1622–1625 (2004). 10.1126/science.1099390

10 Kussell, E., Kishony, R., Balaban, N. Q. & Leibler, S. Bacterial persistence: a model of survival in changing environments. Genetics 169, 1807–1814 (2005). 10.1534/genetics.104.035352

11 Wood, T. K., Knabel, S. J. & Kwan, B. W. Bacterial persister cell formation and dormancy. Appl Environ Microbiol 79, 7116–7121 (2013). 10.1128/AEM.02636-13

12 Oren, Y. et al. Cycling cancer persister cells arise from lineages with distinct programs. Nature 596, 576–582 (2021). 10.1038/s41586-021-03796-6

13 Bressloff, P. C. Stochastic switching in biology: from genotype to phenotype. J. Phys. A: Math. Theor. 50 (2017).

14 Tian, T. & Burrage, K. Stochastic models for regulatory networks of the genetic toggle switch. Proc Natl Acad Sci U S A 103, 8372–8377 (2006). 10.1073/pnas.0507818103

15 Fu, X. et al. Spatial self-organization resolves conflicts between individuality and collective migration. Nat Commun 9, 2177 (2018). 10.1038/s41467-018-04539-4

16 Adler, J. Chemotaxis in bacteria. Science 153, 708–716 (1966). 10.1126/science.153.3737.708

17 Waite, A. J. et al. Non-genetic diversity modulates population performance. Mol Syst Biol 12, 895 (2016). 10.15252/msb.20167044

18 Mattingly, H. H. & Emonet, T. Collective behavior and nongenetic inheritance allow bacterial populations to adapt to changing environments. Proc Natl Acad Sci U S A 119, e2117377119 (2022). 10.1073/pnas.2117377119

19 Wong-Ng, J., Celani, A. & Vergassola, M. Exploring the function of bacterial chemotaxis. Curr Opin Microbiol 45, 16–21 (2018). 10.1016/j.mib.2018.01.010

20 Koster, D. A., Mayo, A., Bren, A. & Alon, U. Surface growth of a motile bacterial population resembles growth in a chemostat. J Mol Biol 424, 180–191 (2012). 10.1016/j.jmb.2012.09.005

21 Cremer, J. et al. Chemotaxis as a navigation strategy to boost range expansion. Nature 575, 658–663 (2019). 10.1038/s41586-019-1733-y

22 Zheng, H. et al. Interrogating the Escherichia coli cell cycle by cell dimension perturbations. Proc Natl Acad Sci U S A 113, 15000–15005 (2016). 10.1073/pnas.1617932114

23 Wolfe, A. J. & Berg, H. C. Migration of bacteria in semisolid agar. Proc Natl Acad Sci U S A 86, 6973–6977 (1989). 10.1073/pnas.86.18.6973

24 Staropoli, J. F. & Alon, U. Computerized analysis of chemotaxis at different stages of bacterial growth. Biophys J 78, 513–519 (2000). 10.1016/S0006-3495(00)76613-6

25 Pleska, M., Jordan, D., Frentz, Z., Xue, B. & Leibler, S. Nongenetic individuality, changeability, and inheritance in bacterial behavior. Proc Natl Acad Sci U S A 118 (2021). 10.1073/pnas.2023322118

26 Vorobyeva, N. V., Sherman, M. & Glagolev, A. N. Bacterial chemotaxis controls the catabolite repression of flagellar biogenesis. FEBS Lett 143, 233–236 (1982). 10.1016/0014-5793(82)80106-3

27 Scott, M. & Hwa, T. Shaping bacterial gene expression by physiological and proteome allocation constraints. Nat Rev Microbiol 21, 327–342 (2023). 10.1038/s41579-022-00818-6

28 Licata, N. A., Mohari, B., Fuqua, C. & Setayeshgar, S. Diffusion of Bacterial Cells in Porous Media. Biophys J 110, 247–257 (2016). 10.1016/j.bpj.2015.09.035

29 Dufour, Y. S., Fu, X., Hernandez-Nunez, L. & Emonet, T. Limits of feedback control in bacterial chemotaxis. PLoS Comput Biol 10, e1003694 (2014). 10.1371/journal.pcbi.1003694

30 Barker, C. S., Pruss, B. M. & Matsumura, P. Increased motility of Escherichia coli by insertion sequence element integration into the regulatory region of the flhD operon. J Bacteriol 186, 7529–7537 (2004). 10.1128/JB.186.22.7529-7537.2004

31 Luo, C. et al. Genome sequencing of environmental Escherichia coli expands understanding of the ecology and speciation of the model bacterial species. Proc Natl Acad Sci U S A 108, 7200–7205 (2011). 10.1073/pnas.1015622108

32 Tu, Y., Shimizu, T. S. & Berg, H. C. Modeling the chemotactic response of Escherichia coli to time-varying stimuli. Proc Natl Acad Sci U S A 105, 14855–14860 (2008). 10.1073/pnas.0807569105

33 Keegstra, J. M. et al. Phenotypic diversity and temporal variability in a bacterial signaling network revealed by single-cell FRET. Elife 6 (2017). 10.7554/eLife.27455

34 Salman, H. & Libchaber, A. A concentration-dependent switch in the bacterial response to temperature. Nat Cell Biol 9, 1098–1100 (2007). 10.1038/ncb1632

35 Yang, Y. et al. Relation between chemotaxis and consumption of amino acids in bacteria. Mol Microbiol 96, 1272–1282 (2015). 10.1111/mmi.13006

36 Li, M. & Hazelbauer, G. L. Cellular stoichiometry of the components of the chemotaxis signaling complex. J Bacteriol 186, 3687–3694 (2004). 10.1128/JB.186.12.3687-3694.2004

37 Blattner, F. R. et al. The complete genome sequence of Escherichia coli K-12. Science 277, 1453–1462 (1997). 10.1126/science.277.5331.1453

38 Adler, J., Hazelbauer, G. L. & Dahl, M. M. Chemotaxis toward sugars in Escherichia coli. J Bacteriol 115, 824–847 (1973). 10.1128/jb.115.3.824-847.1973

39 Parkinson, J. S. & Houts, S. E. Isolation and behavior of Escherichia coli deletion mutants lacking chemotaxis functions. J Bacteriol 151, 106–113 (1982). 10.1128/jb.151.1.106-113.1982

40 Endres, R. G. & Wingreen, N. S. Precise adaptation in bacterial chemotaxis through “assistance neighborhoods”. Proc Natl Acad Sci U S A 103, 13040–13044 (2006). 10.1073/pnas.0603101103

41 Sourjik, V. Receptor clustering and signal processing in E. coli chemotaxis. Trends Microbiol 12, 569–576 (2004). 10.1016/j.tim.2004.10.003

42 Parkinson, J. S. cheA, cheB, and cheC genes of Escherichia coli and their role in chemotaxis. J Bacteriol 126, 758–770 (1976). 10.1128/jb.126.2.758-770.1976

43 Bhattacharjee, T. & Datta, S. S. Bacterial hopping and trapping in porous media. Nat Commun 10, 2075 (2019). 10.1038/s41467-019-10115-1

44 Kurzthaler, C. et al. A geometric criterion for the optimal spreading of active polymers in porous media. Nat Commun 12, 7088 (2021). 10.1038/s41467-021-26942-0

45 Joers, A. & Tenson, T. Growth resumption from stationary phase reveals memory in Escherichia coli cultures. Sci Rep 6, 24055 (2016). 10.1038/srep24055

46 Veening, J. W. et al. Bet-hedging and epigenetic inheritance in bacterial cell development. Proc Natl Acad Sci U S A 105, 4393–4398 (2008). 10.1073/pnas.0700463105

47 Bhattacharyya, S. et al. A heritable iron memory enables decision-making in Escherichia coli. Proc Natl Acad Sci U S A 120, e2309082120 (2023). 10.1073/pnas.2309082120

48 Gude, S. et al. Bacterial coexistence driven by motility and spatial competition. Nature 578, 588–592 (2020). 10.1038/s41586-020-2033-2

49 Goldford, J. E. et al. Emergent simplicity in microbial community assembly. Science 361, 469–474 (2018). 10.1126/science.aat1168

50 Bai, Y. et al. Spatial modulation of individual behaviors enables an ordered structure of diverse phenotypes during bacterial group migration. Elife 10 (2021). 10.7554/eLife.67316

51 Zuo, W. & Wu, Y. Dynamic motility selection drives population segregation in a bacterial swarm. Proc Natl Acad Sci U S A 117, 4693–4700 (2020). 10.1073/pnas.1917789117

52 Venkiteswaran, G. et al. Generation and dynamics of an endogenous, self-generated signaling gradient across a migrating tissue. Cell 155, 674–687 (2013). 10.1016/j.cell.2013.09.046

53 Scherber, C. et al. Epithelial cell guidance by self-generated EGF gradients. Integr Biol (Camb) 4, 259–269 (2012). 10.1039/c2ib00106c

54 David, N. B. et al. Molecular basis of cell migration in the fish lateral line: role of the chemokine receptor CXCR4 and of its ligand, SDF1. Proc Natl Acad Sci U S A 99, 16297–16302 (2002). 10.1073/pnas.252339399

55 Theveneau, E. & Mayor, R. Neural crest delamination and migration: from epithelium-to-mesenchyme transition to collective cell migration. Dev Biol 366, 34–54 (2012). 10.1016/j.ydbio.2011.12.041

56 Muinonen-Martin, A. J. et al. Melanoma cells break down LPA to establish local gradients that drive chemotactic dispersal. PLoS Biol 12, e1001966 (2014). 10.1371/journal.pbio.1001966

57 Acar, M., Mettetal, J. T. & van Oudenaarden, A. Stochastic switching as a survival strategy in fluctuating environments. Nat Genet 40, 471–475 (2008). 10.1038/ng.110

58 Mattingly, H. H., Kamino, K., Machta, B. B. & Emonet, T. Escherichia coli chemotaxis is information limited. Nat Phys 17, 1426–1431 (2021). 10.1038/s41567-021-01380-3

